# A single-nucleotide change in the Kozak sequence enhances protein expression

**DOI:** 10.64898/2026.05.20.726450

**Authors:** Renzo A. Knol, Tahmina Fariaby, Eveline A. de Vlieger, Alexander Kros, Bram Slütter

**Affiliations:** Division of BioTherapeutics, Leiden Academic Centre for Drug Research, Leiden University, Leiden, The Netherlands; Department of Supramolecular & Biomaterials Chemistry, Leiden Institute of Chemistry, Leiden University, Leiden, The Netherlands

**Keywords:** mRNA, *in vitro* transcription, Kozak sequence, translation initiation site, untranslated regions, lipid nanoparticles, gene therapy

## Abstract

*In vitro*-transcribed messenger RNA (IVT mRNA) has emerged as a versatile protein expression platform with broad clinical potential. Current optimization strategies for IVT mRNA focus on untranslated regions (UTRs), mRNA stability, and codon usage, often guided by massively parallel screening and machine learning approaches. In contrast, the Kozak sequence, a key determinant of translation initiation, is often inconsistently incorporated into synthetic 5′ UTR design, and its contribution to translation efficiency remains poorly defined. Here, we systematically varied the Kozak sequence across diverse UTR contexts and performed combinatorial optimization using synthetic, established, and viral UTRs to identify design principles for enhanced translation. We show that a single-nucleotide deviation from the consensus Kozak sequence consistently enhances protein expression across UTR contexts and coding sequences. This effect is conserved across *in vitro* and *in vivo* models, highlighting the generalizability of the optimized Kozak sequence. These findings redefine the role of the Kozak sequence in synthetic mRNA design and demonstrate its substantial contribution to translation efficiency when optimized, enabling improved mRNA-based therapeutics.

## Introduction

Messenger RNA (mRNA) has emerged as a versatile platform for the expression of virtually any protein of interest. Despite sequence- and structure-dependent differences, mRNAs share conserved physicochemical properties as negatively charged polymers composed of four nucleobases and structurally consist of a 5′ cap, 5′ untranslated region (UTR), coding sequence (CDS; open reading frame, ORF), 3′ UTR, and poly(A) tail. Together, these elements regulate translation efficiency and mRNA stability through modulation of ribosome recruitment, translation initiation, and transcript stability.^1–4^ Unlike DNA, which can activate cytosolic DNA-sensing pathways and induce strong innate immune responses,^5^ mRNA is a native cytosolic molecule whose immunogenicity can be modulated by nucleoside modifications,^6,7^ a crucial discovery for harnessing mRNA as a pharmaceutical. Together, these conserved physicochemical and structural characteristics enable efficient, scalable mRNA synthesis, exemplified by the global deployment of mRNA vaccines during the COVID-19 pandemic. ^8,9^

Key foundations for therapeutic mRNA design were established in the 1980s with the identification of the *Kozak consensus sequence* (5′-(gcc)gccRccAUGG-3′), a central determinant of translation initiation^3^, and stabilizing elements within globin UTRs,^10–13^ which later enabled enhanced translation of *in vitro*-transcribed (IVT) mRNA.^14,15^ More recently, unbiased screening approaches identified synthetic and endogenous UTR combinations that further improve IVT mRNA stability and protein expression.^16^ Notably, the AES-mtRNR1 3′ UTR combination outperformed the human β-globin 3′ UTR in multiple systems^16^ and was subsequently incorporated into the Comirnaty vaccine.^17^

Recent efforts to improve translation from IVT mRNA have increasingly relied on massively parallel screening and machine learning approaches to optimize UTRs,^18–22^ mRNA stability,^23^ and codon usage.^24^ These strategies enabled large-scale exploration of sequence–function relationships and led to the identification of synthetic UTRs that outperform commonly used globin-derived UTRs *in vitro* and *in vivo*.^25,26^ In addition, massively parallel reporter assays integrating UTR, codon, and RNA structure analyses identified Dengue virus UTRs among the strongest ribosome recruiters, despite their relatively limited overall translation performance.^27^ Building on these findings, more recent studies explored combinatorial optimization of (synthetic) UTR elements, although these approaches either preselected candidate 5′ UTRs for pairing with established 3′ UTRs^28^ or focused on additional parameters such as cap structure and poly(A) tail length,^29^ thereby limiting systematic exploration of UTR combinations.

As a result, the combinatorial (synthetic) UTR design space remains incompletely explored. In addition, the Kozak sequence is often inconsistently incorporated into synthetic 5′ UTR designs, either appended after UTR generation^26,28^ or variably embedded within sequence optimization workflows.^19,25^ Consequently, subsequent validation studies and combinatorial optimization of UTRs are frequently confounded by variations in the Kozak sequence.^25,29^ Critically, the contribution of the Kozak sequence to translation initiation is only indirectly assessed in massively parallel screening studies, which typically focus on translation initiation site (TIS) features within limited positional windows and primarily recapitulate canonical nucleotide preferences at the key –3 and +4 positions relative to the start codon.^19,21^

Importantly, although the canonical vertebrate Kozak consensus sequence is widely implemented in transgene and IVT mRNA design, its degree of conservation among endogenous vertebrate transcripts is remarkably low. Early analyses estimated that only ∼0.2% of vertebrate mRNAs perfectly match the full GCCGCCACC consensus sequence.^30^ Moreover, naturally occurring sequence variation in Kozak regions has repeatedly been linked to altered translation efficiency^31^ and human disease.^32–35^ Comparative analyses have further identified species-specific Kozak preferences that significantly influence translation efficiency *in vivo*. In zebrafish embryos, small nucleotide differences relative to the canonical vertebrate consensus were shown to substantially enhance protein expression.^36^ Together, these findings suggest that the canonical Kozak sequence may not represent the globally optimal architecture in all biological or therapeutic contexts, and that the term *consensus* should not be regarded as optimal.^30^

However, despite previous efforts to derive alternative Kozak architectures from tissue-specific or species-specific transcript datasets,^36,37^ the Kozak sequence has rarely been systematically evaluated as an independent design element across multiple IVT mRNA contexts. In particular, its contribution to translation efficiency relative to surrounding UTR architectures remains insufficiently characterized in IVT mRNA systems and has not been broadly validated across *in vitro* and *in vivo* delivery settings.

Here, we systematically assessed minimally modified Kozak sequence variants across multiple 5′ UTR contexts to identify determinants of enhanced translation from IVT mRNA. We subsequently performed combinatorial optimization of previously reported promising (synthetic) UTRs *in vitro* and in zebrafish embryos, followed by validation of top-performing combinations across additional Kozak variants and in mice. Notably, a single-nucleotide deviation from the consensus Kozak sequence consistently enhanced protein expression across UTRs, coding sequences, and model systems. The two best-performing combinations were further tested in a therapeutic setting in low-density lipoprotein receptor-deficient (*Ldlr ^-/-^*) mice, where they outperformed the Comirnaty UTRs in restoring Ldlr expression and reducing serum cholesterol levels. Together, these findings establish Kozak sequence design as a major determinant of translation efficiency and support the rational design of more potent mRNA therapeutics.

## Results

### Combinatorial UTR selection and Kozak sequence variation design

When assessing candidate UTRs generated or evaluated in previous studies, we noticed that the Kozak sequence (5′-(gcc)gccRccAUGG-3′) design was inconsistent, with studies appending either a minimal (GCCACC)^28^ or an extended (GCCGCCACC)^26^ consensus Kozak sequence variant after generating synthetic 5′ UTRs. In contrast, another strategy directly incorporated the Kozak sequence into the 5′ UTR generation process.^19^ Selected top-performing synthetic UTRs selected from previous studies were the 5′ UTRs NeoUTR3^26^ (hereafter named Syn1) and UTR4^19^ (hereafter named Syn2). Additionally, the highly stabilizing 3′ UTR mtRNR1 and the AES-mtRNR1^16^ combination were included. The Dengue virus-derived UTRs were included as they were previously identified among the best ribosome recruiters but failed to induce high protein expression,^27^ likely due to the lack of a Kozak sequence. Human β-globin UTRs (HBB) were included as golden standards, supplemented with the Comirnaty UTR combination (human α-globin 5′ UTR with AES-mtRNR1 3′ UTR, hereafter abbreviated as BNT).^17^ Importantly, the selected 5′ UTRs have inconsistent Kozak sequences, which likely significantly impacts UTR performance (Figure 1A).

**Figure 1.**
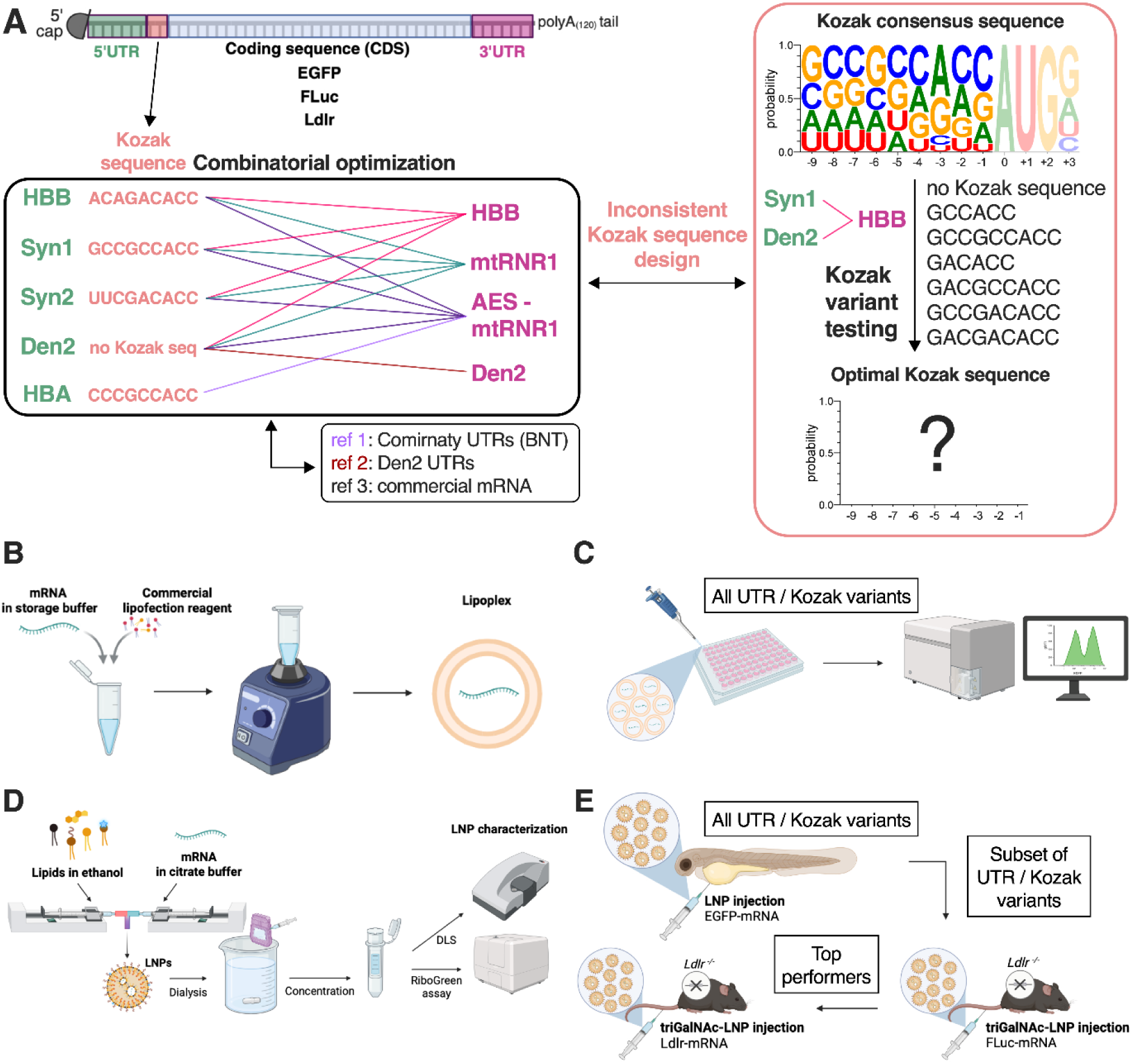
Schematic representation of the experimental design and procedures used in this study. **A)** *In vitro*-transcribed mRNAs carrying various combinations of previously reported (Syn1 is NeoUTR3^26^, Syn2 is UTR4^25^) untranslated regions (UTRs), along with several references, are selected for combinatorial optimization. Kozak sequence design is inconsistent across these reported UTRs. Notable variations include substitutions of guanosine to adenosine at the –5 position relative to the start codon (in HBB and Syn2). G-to-A substitutions at the –5 and/or –7 positions are therefore tested in expression screening studies to identify an optimal Kozak sequence. **B)** All selected Kozak variants and UTR combinations are evaluated with the EGFP coding sequence. EGFP-mRNA is complexed with a commercial lipofection reagent. **C)** Lipoplexes are transfected into four different cell lines, and EGFP expression levels are measured 24 h later by flow cytometry. **D)** Encapsulation of mRNAs in lipid nanoparticles (LNPs) for *in vivo* expression analysis is performed by microfluidic mixing, dialysis against PBS, centrifugation-based concentration, and subsequent physicochemical characterization. **E)** All selected Kozak variants and UTR combinations are evaluated in zebrafish embryos. A subset is selected for screening in low-density lipoprotein receptor-deficient (*Ldlr ^-/-^*) mice using firefly luciferase (FLuc) encapsulated in triantennary N-acetylgalactosamine (triGalNAc)-functionalized LNPs. Top performers are evaluated in a therapeutic setting by comparing their ability to temporarily restore Ldlr expression in the liver, thereby reducing serum cholesterol levels.

When critically assessing the Kozak sequences of the selected 5′ UTRs, we noticed that HBB and Syn2 both bear an adenosine rather than a guanosine at the –5 position relative to the start codon. Since both 5′ UTRs are among the most potent translation initiators to date, we reasoned that minimally varying the consensus Kozak sequence with adenosines at the –5 and/or –7 positions might enhance translation initiation efficiency. Additionally, previous species- and tissue-specific advantages were found for an adenosine at the –5 position.^37,38^ Therefore, we selected the Syn1 and Den2 5′ UTRs, originally reported with and without the Kozak consensus sequence, respectively, for systematic testing of Kozak sequence variants (Figure 1A). Other factors in mRNA design, such as cap structure, nucleoside modification, a template-encoded poly(A_120_) tail, the EGFP coding sequence, and the HBB 3′ UTR, were kept constant.

### An adenosine at the –5 position of the Kozak sequence increases protein expression

To assess potential Kozak sequence-dependent EGFP expression differences across *in vitro* model systems, EGFP-mRNAs were complexed with a commercial lipofection reagent for transfection into four cell lines (Figure 1B): the murine dendritic cell line DC2.4, human umbilical vein endothelial cells (HUVECs), the human embryonic kidney cell line HEK293T, and the human cervical cancer cell line HeLa. Interestingly, flow cytometry revealed that the non-canonical Kozak sequence GCCGACACC yielded the highest EGFP expression, regardless of the cell line or 5′ UTR used. The only exception was the GACACC Kozak variant that was superior in one specific case; being for the Syn1 5′ UTR in HUVECs. Importantly, a guanosine to adenosine replacement on the –7 position was not beneficial, and adenosines on both –5 and –7 positions further reduced EGFP expression (Figure 2; Figure S1). Finally, as expected, the absence of a Kozak sequence was detrimental to EGFP expression, although Den2 performed better without a Kozak sequence than Syn1 (Figure S1). Collectively, these results suggest that an adenosine at the –5 position relative to the start codon improves protein expression *in vitro*.

**Figure 2.**
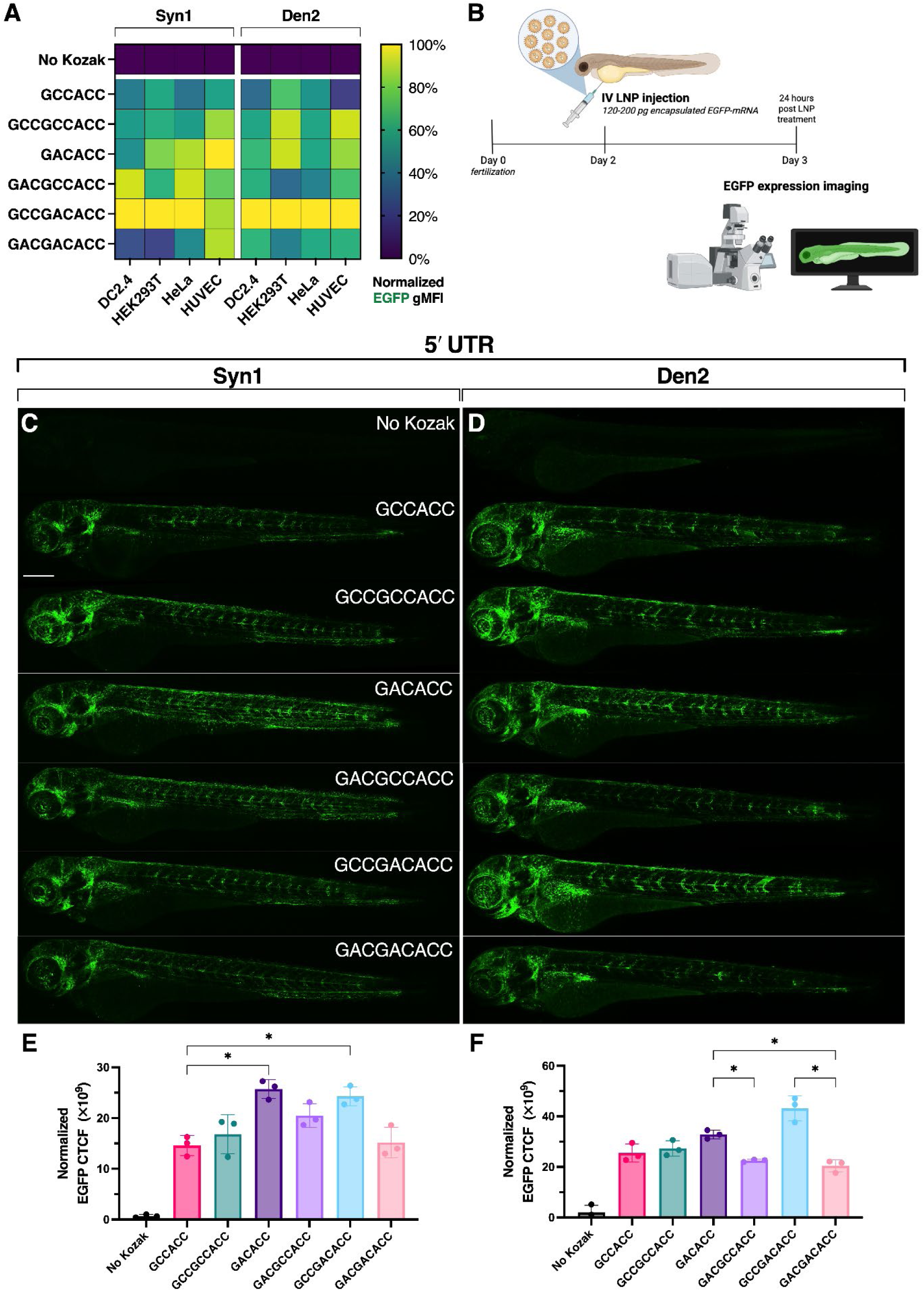
Effect of Kozak sequence variation on protein expression *in vitro* and in zebrafish embryos. **A)** Heatmap showing relative EGFP expression levels, normalized to the highest group per cell line, 24 h after lipofection with 500 ng mL^-1^ EGFP-mRNA carrying Kozak sequence variants of either the Syn1 or Den2 5′ UTR paired with the HBB 3′ UTR. **B)** Schematic of the general experimental procedure for expression analysis in zebrafish embryos. Embryos received 200 pg (0.2 mg/kg) EGFP-mRNA encapsulated in LNPs. 24 h later, whole-body EGFP expression was imaged by confocal microscopy. **C, D)** Confocal z-projections showing EGFP expression (green) in representative wild-type (AB/TL) zebrafish embryos 24 h after receiving LNP-encapsulated (i.v.) 0.2 mg/kg EGFP-mRNA carrying either Kozak variant and the Syn1 (**C**) or Den2 (**D**) 5′ UTR paired with the HBB 3′ UTR. Scale bar represents 250 µm. **E, F)** Quantification of EGFP expression levels in wild-type zebrafish embryos that received Kozak sequence variants with Syn1 (**E**) or Den2 (**F**) 5′ UTRs by corrected total cell fluorescence (CTCF). Data are presented as group mean (n=3) ± SD and statistically compared using a Brown-Forsythe & Welch ANOVA with Dunnett T3 correction (* p>0.05).

Encouraged by these *in vitro* findings, we turned to zebrafish embryos, a convenient, medium- to high-throughput screening model that enables live nanoparticle biodistribution and protein expression analysis.^39–44^ Two-day-old embryos intravenously received 0.2 mg/kg EGFP-mRNA encapsulated in lipid nanoparticles (LNPs) (Figure 2B; Figure S2; Table S4) bearing either the Kozak sequence variant paired with the Syn1 or Den2 5′ UTR. 24 hours later, confocal imaging revealed Kozak sequence-dependent EGFP expression differences: Syn1 performed best with the GACACC variant, whereas Den2 yielded the highest expression with GCCGACACC. Importantly, the next best Kozak variant for each 5′ UTR also carried an adenosine at the –5 position but not at the –7 position (Figure 2C–F). Remarkably, EGFP expression patterns in zebrafish largely match the *in vitro* results, particularly those observed for HUVECs, although a minimal GCCACC Kozak sequence performed relatively better in zebrafish embryos, and overall differences between Kozak variant groups were smaller (Figure S2D).

### Den2 5′ UTR paired with the HBB 3′ UTR enhance EGFP expression in vitro

Next, we performed combinatorial optimization using previously reported UTRs. The UTR combinations were initially evaluated with their original Kozak sequences (Figure 1A), except for the Den2 5′ UTR, which lacks a Kozak sequence. Therefore, as with Syn1, the consensus Kozak sequence (GCCGCCACC) was appended to the Den2 5′ UTR. To compare these UTR designs against a commercial benchmark, TriLink 5-methoxyuridine (5-moU)-modified EGFP-mRNA was included. The resulting EGFP-mRNAs received the same CleanCap® AG cap, a PCR-template-encoded poly(A_120_) tail, the same 5-moU modification, and, to reduce dsRNA byproducts,^45^ a mutant T7 polymerase operating at 50 °C was employed.^46^ To assess their performance across different *in vitro* model systems, EGFP-mRNAs with various UTR combinations were tested in the same cell lines used for Kozak variant screening. To gauge the relative contribution of the Kozak sequence to protein expression, two mRNAs lacking a 5′ UTR but carrying the minimal GCCACC or optimal GCCGACACC Kozak sequences paired with the HBB 3′ UTR were included.

Interestingly, the Den2 5′ UTR paired with the HBB 3′ UTR was consistently superior across cell lines. The synthetic 5′ UTRs paired with HBB performed nearly as well in HEK293T, and Syn2 performed well in HeLa. Among the 3′ UTRs, mtRNR1 performed the worst overall, but performance within that group was strongly dependent on the 5′ UTR and cell line. In contrast, within the AES-mtRNR1 group, expression levels across 5′ UTRs and cell lines varied the least, highlighting its general stabilizing effect. The BNT UTRs performed at an intermediate level across cell lines, and the Den2 5′ UTR performed worse when paired with its matching 3′ UTR than with the other 3′ UTRs. Moreover, mRNAs lacking 5′ UTRs showed intermediate expression levels, with the GCCGACACC variant clearly outperforming the minimal GCCACC variant, at levels similar to those of the BNT UTRs across cell lines. Although the commercial EGFP-mRNA was superior in HEK293T, it ranked second in DC2.4 and HUVECs and lower in HeLa (Figure 3A; Figure S3). Collectively, these results identified the Den2 5′ UTR paired with HBB, despite the suboptimal Kozak sequence used, as the optimal tested UTR combination *in vitro*.

**Figure 3.**
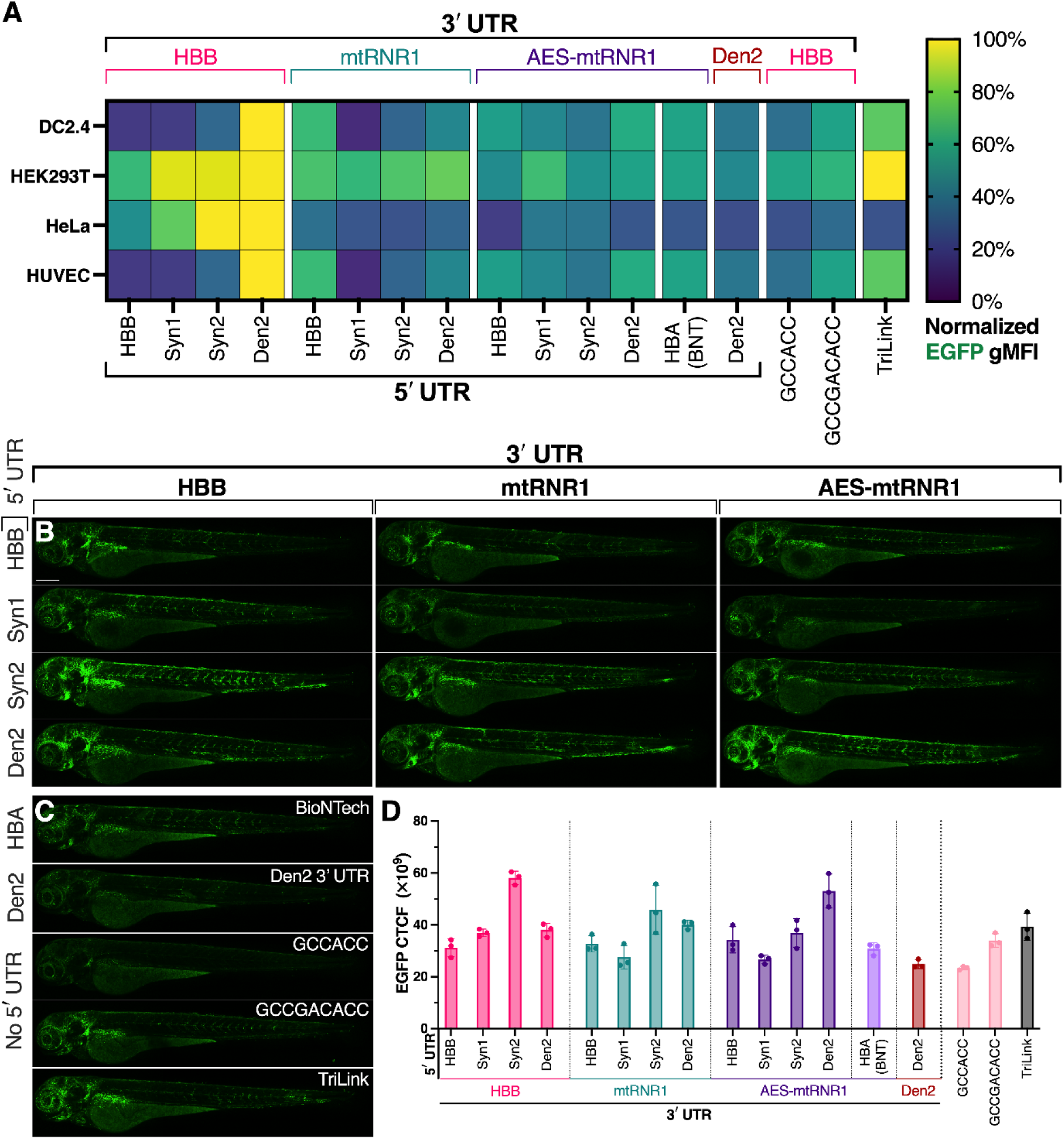
Combinatorial UTR optimization *in vitro* and in zebrafish embryos. **A)** Heatmap depicting relative EGFP expression levels normalized to the highest group per cell line 24 h after lipofection with 500 ng mL^-1^ EGFP-mRNAs carrying various UTR combinations. **B, C)** Confocal z-projections showing EGFP expression (green) of representative wild-type (AB/TL) zebrafish embryos 24 h after receiving (i.v.) 0.2 mg/kg EGFP-mRNA carrying different UTR combinations (**B**) or references (**C**) encapsulated in Onpattro-like LNPs. Scale bar represents 250 µm. **D)** Quantification of EGFP expression levels in wild-type zebrafish embryos that received EGFP-mRNA carrying different UTR combinations by corrected total cell fluorescence (CTCF).

### Syn2 5′ UTR combined with HBB 3′ UTR yields high EGFP expression in zebrafish embryos

To assess whether the *in vitro* findings translate to an *in vivo* model, the same EGFP-mRNAs with various UTR combinations, along with the references, were encapsulated in LNPs (Figure 1D–E; Table S4) and intravenously administered to two-day-old zebrafish embryos. 24 hours later, confocal imaging showed that Syn2 paired with the HBB 3′ UTR performed best. In fact, Syn2 was outperformed only by Den2 when paired with AES-mtRNR1, and across 3′ UTRs, the Den2 5′ UTR ranked second after Syn2. In contrast, Syn1 performed among the worst regardless of the paired 3′ UTR. Moreover, consistent with the *in vitro* results, BNT UTRs performed at an intermediate level, similar to the GCCGACACC–HBB variant without a 5′ UTR (Figure 3B–D; Figure S4). Collectively, 5′ UTRs carrying an adenosine at the –5 position generally performed better than those containing a guanosine at that position, regardless of paired 3′ UTR, except for Den2, which generally performed well.

### Optimal Kozak sequence integration enhances protein expression across UTRs

To further confirm the previous findings, A-to-G substitutions at the –5 position were introduced into HBB and Syn2, and the resulting sequences were compared with their original sequences. Additionally, a stronger focus was placed on the Den2 5′ UTR due to its overall high performance; therefore, Den2 was paired with the mtRNR1 and AES-mtRNR1 3′ UTRs, using the optimized Kozak sequence (GCCGACACC). To confirm whether truncated versions of the Den2 5′ UTR would provide no further benefit to protein output with the optimized Kozak sequence (GCCGACACC), 50-nt and 25-nt truncated versions were generated with the optimized Kozak sequence appended and paired with HBB 3′ UTR. Additionally, because Syn2 also performed well in previous experiments, a similar approach was taken, yielding a 25-nt truncated version.

Importantly, suboptimal Kozak sequences in HBB and Syn2 led to reduced EGFP expression *in vitro* (Figure 4A). This effect was even more pronounced in zebrafish embryos, where the original Syn2 Kozak sequence ((UUC)GACACC) outperformed the best-performing *in vitro* Kozak sequence (GCCGACACC) (Figure 4B, D; Figure S5). As previously observed, Den2, when paired with mtRNR1 or AES-mtRNR1, yielded lower protein expression *in vitro* than with HBB.

**Figure 4.**
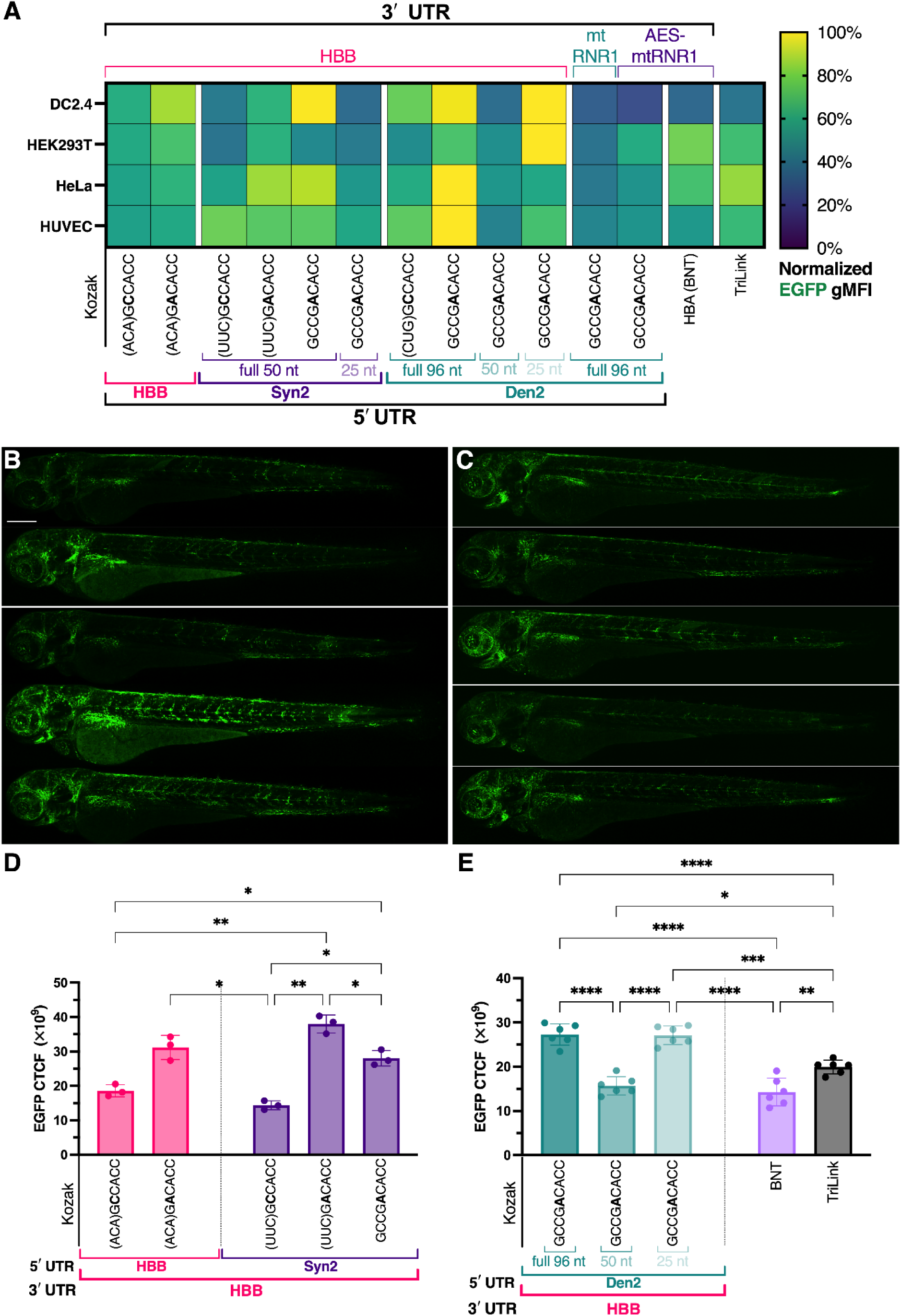
Combined effect of a G-to-A substitution at the –5 position and 5′ UTR truncation on EGFP expression. **A)** Heatmap showing relative EGFP expression levels, normalized to the highest group per cell line, 24 h after lipofection with 500 ng mL^-1^ EGFP-mRNAs carrying different Kozak sequences and (truncated) UTR variants. **B)** Confocal z-projections showing EGFP expression (green) in representative wild-type (AB/TL) zebrafish embryos 24 h after receiving (i.v.) 0.2 mg/kg EGFP-mRNA carrying HBB or Syn2 with different Kozak variants encapsulated in LNPs. Scale bar represents 250 µm. **C)** Confocal z-projections showing EGFP expression in representative wild-type zebrafish embryos 24 h after receiving (i.v.) 0.12 mg/kg EGFP-mRNA carrying full-length or truncated Den2 5′ UTR variants or reference EGFP-mRNAs encapsulated in DiD-labeled LNPs. **D, E)** Quantification of EGFP expression levels in wild-type zebrafish embryos that received Kozak sequence variants with HBB or Syn2 (**D**) or full-length or truncated Den2 5′ UTRs (**E**) by corrected total cell fluorescence (CTCF). Data are presented as group means (n=3 or 6) ± SD and are statistically compared by a Brown-Forsythe & Welch ANOVA with Dunnett T3 correction (**D**) or a regular one-way ANOVA with Tukey correction (**E**) (* p>0.05, ** p>0.01, *** p>0.001, **** p>0.0001).

Translation efficiency is strongly influenced by 5′ UTR length and composition, as longer UTRs increase the likelihood of inhibitory features, including secondary structure, upstream open reading frames (uORFs), and regulatory elements.^2,47–49^ This suggests that simpler, shorter 5′ UTRs might be beneficial. Moreover, shorter 5′ UTRs can simplify PCR-based IVT template generation, as they are more readily incorporated into primer overhangs, facilitating rapid and scalable mRNA production workflows. Particularly, Den2 has been reported to contain strong secondary structures^27^ that could reduce accessibility to the TIS.

Although secondary structures in the 5′ UTR can be altered by the CDS or 3′ UTR, RNAfold structure prediction shows that Syn2’s structure is slightly altered when changing the 3′ UTR (Figure S10A). In contrast, the Den2 5′ UTR has consistent stem–loop structures regardless of CDS or 3′ UTR (Figure S10B–C), suggesting a strong dependence on eukaryotic initiation factor proteins to unwind these structures, enabling translation initiation.^49^ Importantly, mRNA structure prediction suggests that removing downstream regions of the 96-nucleotide-long Den2 5′ UTR, leaving only 50 nucleotides (nt) or 25 nt closest to the start codon, also removes the strong stem-loop structure (Figure S10B–C).

Therefore, truncated variants of Syn2 and Den2 containing the optimal Kozak sequence paired with the HBB 3′ UTR were evaluated. *In vitro*, the 25-nt Den2 5′ UTR slightly outperformed its full-length counterpart in DC2.4 cells and showed a more pronounced improvement in HEK293T cells, surpassing the commercial EGFP-mRNA (Figure 4A). In contrast, truncating Syn2 to 25 nt reduced EGFP expression across cell lines. Results in zebrafish embryos for the 50- and 25-nt Den2 variants were in line with findings for DC2.4, with the 50-nt variant showing reduced expression, while the 25-nt Den2 performed similarly to its full-length counterpart (Figure 4C, E). These results further strengthen previous findings on the G-to-A (–5) protein expression improvement and generally imply that truncating the 5′ UTRs provides no benefit.

Finally, three other Kozak variants were tested in the context of the Den2 5′ UTR paired with the HBB 3′ UTR, comparing the sequence identified here as optimal (GCCGACACC) with the previously reported optimized Kozak sequence for retinal genes (ACCGAGACC).^37^ The two additional variants differed from the optimized Kozak sequence only at the –9 position, with A (as in the retinal Kozak) or C (as in the HBA 5′ UTR). These variants were compared with previously screened variants in DC2.4 and HUVECs, as well as in zebrafish embryos. *In vitro*, differences among the –9 position variants were negligible, but they performed substantially worse in zebrafish embryos than GCCGACACC (Figure S8). Conversely, the Kozak sequence optimized for retinal genes performed worse *in vitro* but similarly to the optimal Kozak in zebrafish embryos (Figure S8). Collectively, these results suggest no further improvement by varying the –9 position and no consistent benefit of the retinal Kozak sequence over the optimized Kozak sequence.

### The Den2 5′ UTR yields high luciferase expression in mice

Building on these findings, we sought to investigate a subset of UTR combinations in a mammalian model. Emphasis was placed on the Den2 5′ UTR, which consistently performed well when appended with an (optimized) Kozak sequence. In addition to the full-length Den2 5′ UTR paired with HBB or AES-mtRNR, truncated versions were included. The synthetic 5′ UTRs using the optimized Kozak sequence (GCCGACACC) were paired with HBB. To gauge the relative contribution of the Kozak sequence, only GCCGACACC (lacking a 5′ UTR) was used with the HBB 3′ UTR. The BNT UTRs were included as a reference. The coding sequence was a previously reported red-shifted firefly luciferase,^50^ codon-optimized for mice using an effective high-GC-based strategy from Genewiz, evaluated in a previous study.^27^

With a therapeutic (cholesterol lowering) setting in mind, low-density lipoprotein-deficient (*Ldlr ^-/-^*) mice were used in this experiment. The lack of Ldlr has previously been shown to significantly reduce hepatic targeting due to the absence of the apolipoprotein E (ApoE)–Ldlr hepatic targeting mechanism. An alternative hepatic targeting route uses triantennary N-acetylgalactosamine (triGalNAc) to target the asialoglycoprotein receptor (ASGPR).^51,52^ Therefore, FLuc-mRNA variants were encapsulated using previously optimized triGalNAc-functionalized SM-102-LNPs (hereafter referred to as triGalNAc-LNPs)^52^ (Table S4).

*Ldlr ^-/-^* mice received a 0.15 mg/kg dose of FLuc-mRNA encapsulated in triGalNAc-LNPs via intravenous injection (Figure 5A). 6 hours later, live bioluminescence imaging revealed that, consistent with *in vitro* and zebrafish data, pairing Den2 with the optimized Kozak sequence yielded considerably higher (3.6-fold) FLuc expression than Den2 with GCCACC. As in previous results, the 25-nt Den2 variant, but not the 50-nt variant, produced expression levels similar to those of the full-length variant. Furthermore, Syn1 with the optimal Kozak sequence appended, performed slightly better than Syn2. Interestingly, the variant lacking a 5′ UTR, containing only GCCGACACC, performed remarkably well, nearly matching Den2’s bioluminescence level (Figure 5B, D; Figure S9).

**Figure 5.**
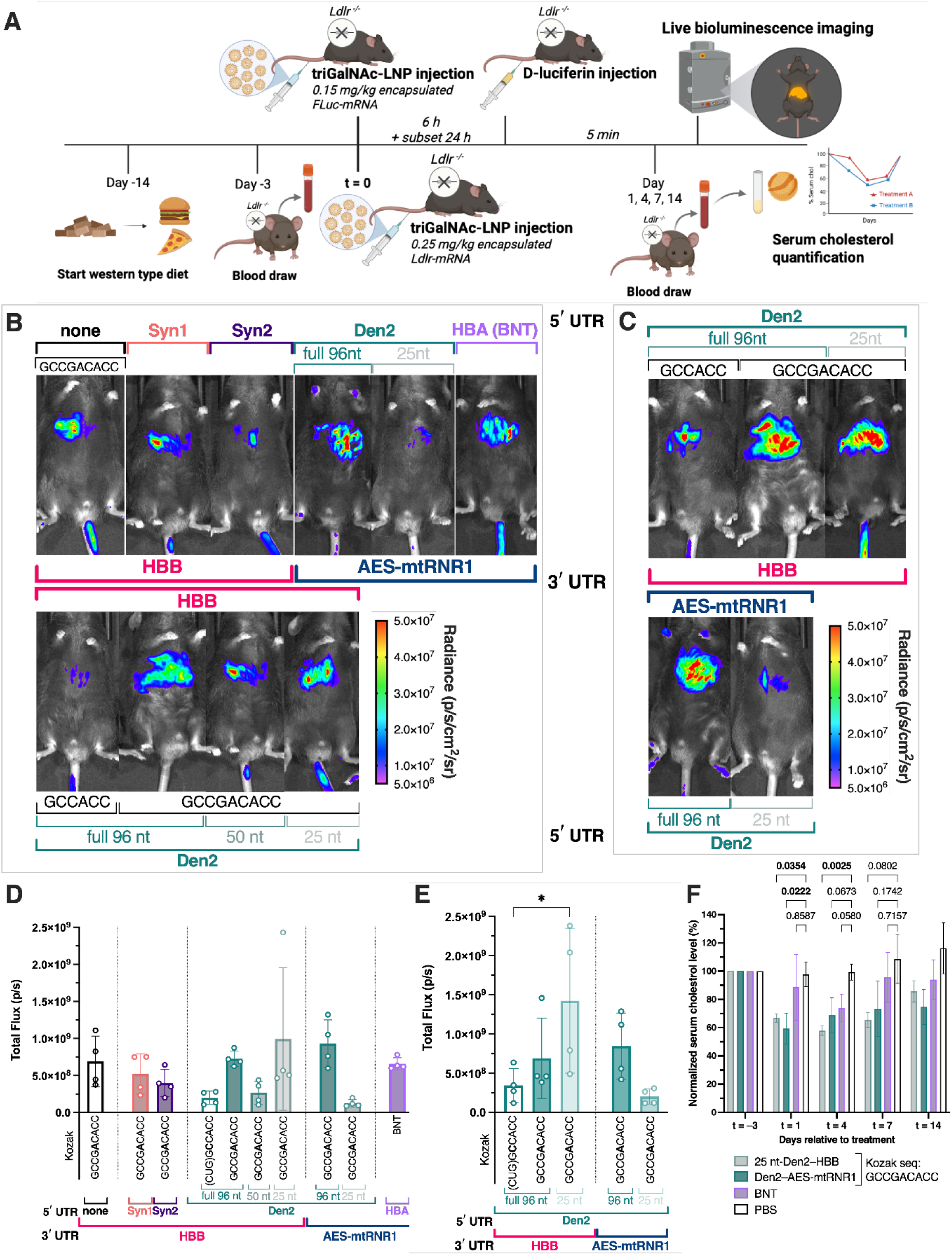
Influence of different UTR or Kozak sequence combinations on protein expression in mice. **A)** Schematic representation of the experimental procedure. Low-density lipoprotein receptor-deficient (*Ldlr ^-/-^*) mice received (i.v.) 0.15 mg/kg FLuc-mRNA or 0.25 mg/kg Ldlr-mRNA encapsulated in triantennary N-acetylgalactosamine (triGalNAc)-functionalized SM-102-LNPs. For bioluminescence measurements, 6 hours later, a second i.v. injection delivered D-luciferin, enabling live bioluminescence imaging reflecting luciferase expression in mice after 5 minutes. A subset of mice was reimaged 5 minutes after a second D-luciferin injection, 24 h after LNP administration. For Ldlr replacement, low-density lipoprotein receptor-deficient (*Ldlr ^-/-^*) mice were fed a Western-type diet starting 14 days before treatment. 3 days before treatment, blood was drawn to establish baseline serum cholesterol levels, allowing normalization to baseline for each individual. After treatment, blood was drawn on days 1, 4, 7, and 14 to measure serum cholesterol levels, which reflect Ldlr activity over time. **B, C)** Live bioluminescence IVIS images of representative individuals at 6 h (**B**) or 24 h (**C**) after receiving 0.15 mg/kg FLuc-mRNA encapsulated in triGalNAc-LNPs carrying different UTR or Kozak sequences. **D, E)** Quantification of FLuc expression by total flux per second at 6 h (**D**) or 24 h (**E**) after LNP administration. Data are presented as group means (n=4) ± SD and statistically compared by one-way ANOVA with Tukey correction (* p>0.05). **F)** Serum cholesterol levels of *Ldlr* ^-*/-*^ mice over time after receiving Ldlr-mRNAs with different UTR variants. Bar plots of the data in panel B showing statistical comparisons of each treatment group with the PBS control over time by two-way ANOVA. Data are presented as group means (n=3) ± SD. Relevant p-values are shown for days 1, 4, and 7 after treatment.

For Den2 with AES-mtRNR1, unlike with HBB, the truncated 25-nt Den2 performed much worse than the full-length variant. mRNA structure prediction suggests that this might be due to reduced accessibility of the translation initiation site (Figure S10C). Moreover, in contrast to *in vitro* data but in line with the zebrafish results, Den2 with AES-mtRNR1 performed slightly better than with HBB. Additionally, the BNT UTRs performed relatively better than *in vitro* or in zebrafish, slightly outperforming the synthetic UTRs (Figure 5B, D; Figure S9).

The initial 6 h bioluminescence measurement was selected based on previous studies indicating peak luciferase expression at 4–6 h after LNP administration.^53–56^ However, the formulation used here employs non-dissociating lipid anchors with functionalized polyethylene glycol (PEG) moieties, which may alter uptake kinetics. To assess temporal effects, mice were reimaged at 24 h post-injection, focusing on the Den2 variants. Expression levels were comparable or higher than at 6 h, with the GCCACC and the 25-nt variants showing elevated expression.

Importantly, in the HBB 3′ UTR group, the 25-nt Den2 exhibited substantially higher bioluminescence in two individual mice, making it the overall best-performing variant. In the AES-mtRNR1 group, expression slightly improved for the low-performing truncated Den2 but remained clearly outperformed by its full-length counterpart (Figure 5C, E; Figure S9). Overall, these results are largely consistent with *in vitro* and zebrafish results despite differences in coding sequence and delivery vehicle, supporting the leading performance of the (truncated) Den2 5′ UTR and the optimized Kozak sequence.

### Den2-Ldlr-mRNA variants with optimal Kozak sequence effectively restore Ldlr expression

To further evaluate the performance of Den2 UTR variants and the optimized Kozak sequence, and to solidify their potency across coding sequences, we sought to perform low-density lipoprotein receptor (Ldlr) replacement therapy in a deficient (*Ldlr ^-/-^*) mouse model. Therefore, the best-performing candidates, 25 nt-Den2 with HBB and full-length Den2 with AES-mtRNR1, were selected. Additionally, the BNT UTRs were included as a clinically approved reference. The native mouse Ldlr coding sequence was codon-optimized for mice using the same strategy previously used for FLuc.

Regarding the experimental procedure, *Ldlr ^-/-^* mice were fed a Western-type diet starting 14 days before treatment to elevate serum cholesterol levels. Three days before treatment, mice were bled to measure cholesterol levels, allowing normalization of cholesterol levels to baseline for each individual throughout the experiment. Mice then received a single 0.25 mg/kg dose of Ldlr-mRNA, either carrying Den2 UTR variants with the optimized Kozak sequence or BNT UTRs, encapsulated in triGalNAc-LNPs. Subsequently, mice were bled on days 1, 4, 7, and 14 after LNP administration to measure serum cholesterol levels, which reflect Ldlr activity over time (Figure 5A).

Ldlr replacement therapy reduced serum cholesterol to ∼58% of the pre-treatment level. Notably, consistent with luciferase expression measurements, the 25 nt-Den2 with HBB combination was most effective overall, reducing serum cholesterol by 33% or more over 7 days, with minimal inter-individual variability. The full-length Den2 with AES-mtRNR1 followed, and the BNT UTRs were least effective. Importantly, the expression dynamics mirrored those observed for FLuc, with the 25 nt-Den2 variant reaching peak expression later than the Den2– AES-mtRNR1 combination (Figure 5F). Collectively, these results support previous findings that Den2 with the optimal Kozak sequence yields the highest protein expression, regardless of the coding sequence or model system.

### An adenosine at the –5 position of the Kozak sequence is optimal across model systems

To comprehensively analyze the generated protein expression data across model systems and identify an overall optimal Kozak sequence, experimentally tested Kozak variants were ranked by relative protein expression within each model system. Expression levels were normalized within each 5′ UTR context to the highest value (Table S5). Subsequently, Kozak sequences were converted into weighted FASTA files, used to visualize the probability of each nucleotide at positions –9 to –1 relative to the start codon.

The Kozak consensus sequences for each model system were extracted from the Ensembl database, revealing minimal differences between the human and mouse sequences. However, the zebrafish start codon context showed a higher probability of adenosines at positions –9 to –7 and –4 to –2. At the –5 position, adenosine probabilities were not among the two highest for any of the model systems. In contrast, combined human cell line data and combined *in vivo* data clearly indicated the highest probability of an adenosine at the –5 position. Differences between combined human cell line data and combined *in vivo* data were minimal, differing mainly in the second- to fourth-highest nucleotide probabilities at positions –9 to –7 (Figure 6A). When all protein expression data were combined, the visualization of nucleotide probabilities clearly indicated that GCCGACACC is the optimal sequence (Figure 6B, C).

**Figure 6.**
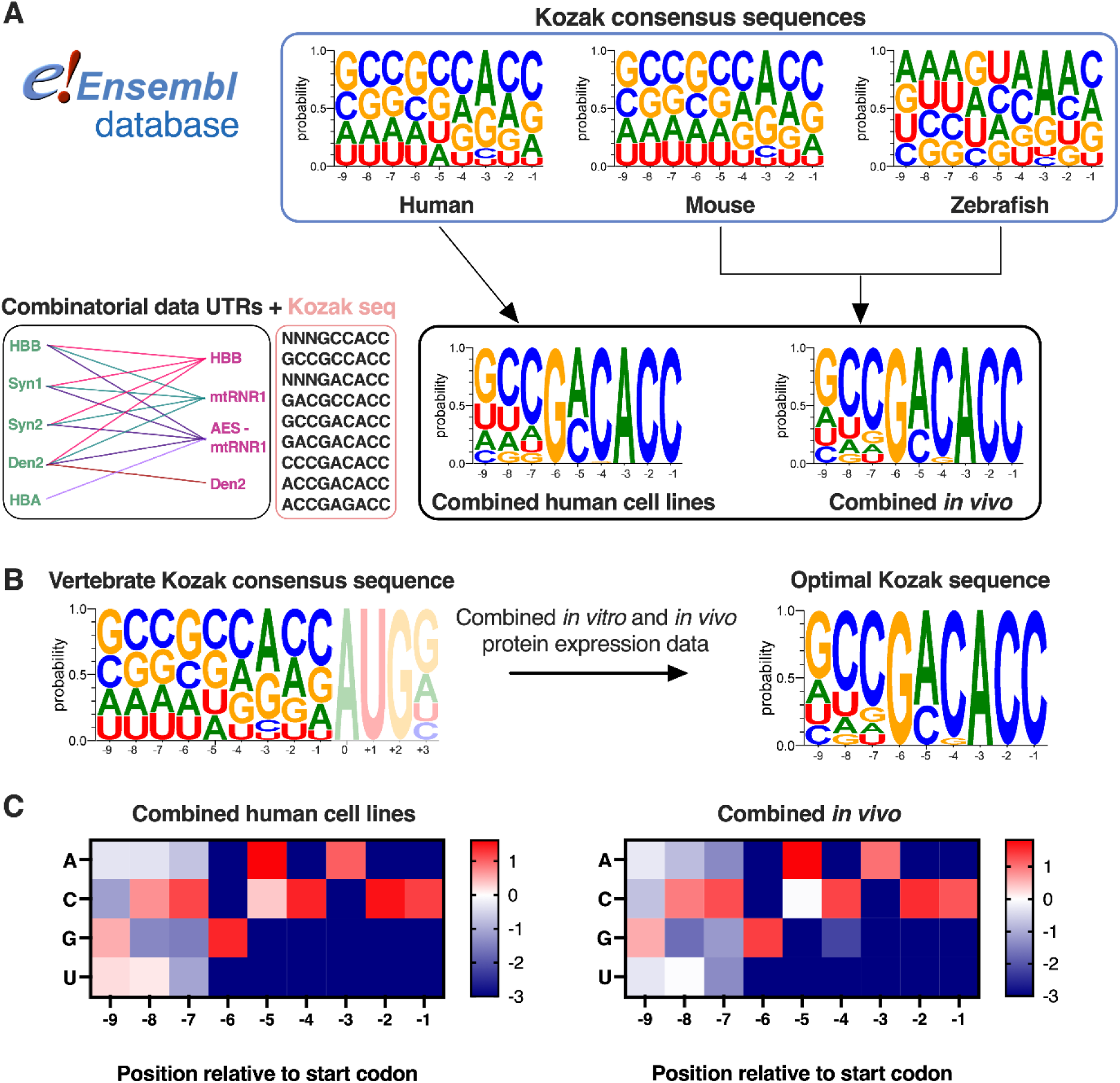
Visualization of Kozak consensus sequences and optimal Kozak sequences based on combined protein expression data across model systems. **A)** Human, mouse, and zebrafish Kozak consensus sequences were extracted from the Ensembl database. Ranked and normalized protein expression data per UTR context in different model systems generated a weighted Kozak sequence list used to visualize nucleotide probabilities on positions –9 to –1 relative to the start codon. **B)** Pooled Ensembl database sequences from human, mouse, and zebrafish generated a vertebrate Kozak consensus sequence. The combined ranked protein expression data across model systems generated a visualization of the generally optimal Kozak sequence. **C)** Enrichment matrices reflecting which nucleotides are enriched at a given position (–9 to –1) in the optimal Kozak sequence compared to the Kozak consensus sequence. Separate matrices were generated for ranked protein expression data from human cell lines and for combined *in vivo* protein expression data. When a nucleotide was absent at a given position, enrichment was capped at −3.

## Discussion

Here, we demonstrate a substantial effect of a G-to-A substitution at the −5 position of the Kozak consensus sequence on protein expression from IVT mRNA. This effect was consistent across various UTR combinations, coding sequences, and model systems, including *in vitro* (Figure 2A, 4A; Figure S1, S3), in zebrafish embryos with EGFP (Figure 2C–F, 4B–E; Figure S2, S4), and in mice with FLuc (Figures 5B–E, S9). Integration of all protein expression data identified an optimal Kozak sequence, GCCGACACC (Figure 6). In a therapeutic setting, Ldlr replacement in *Ldlr ^-/-^* mice further supported the performance of the optimized Kozak sequence (Figure 5F). Although *in vivo* validation was limited to a subset of UTR combinations and liver expression, these findings provide initial evidence for its applicability in a therapeutic context.

The vertebrate Kozak consensus sequence has historically been regarded as a highly efficient translation initiator.^57^ However, several observations in previous literature support the plausibility of alternative variants. The full GCCGCCACC motif is present in only a small fraction of vertebrate transcripts, and nucleotide usage surrounding the start codon differs substantially across eukaryotic species.^30^ Interestingly, enrichment of adenine at the −5 position has been reported in several eukaryotic systems, including yeast, dicot plants, Dictyostelium, and protozoa.^30^ In yeast, experimental studies further demonstrated that the introduction of adenine at −5 can improve translation efficiency.^38^ In addition, previous work identified a species-specific Kozak consensus sequence in zebrafish embryos,^36^ which our findings confirm (Figure 6A), further emphasizing that relatively small nucleotide differences can substantially influence expression outcomes *in vivo*. Importantly, tissue-specific optimization for retinal genes identified an enhanced sequence containing an adenine at the –5 position.^37^ Our findings extend these observations by demonstrating that G-to-A substitution at position −5 enhances translation from therapeutic IVT mRNA constructs across multiple vertebrate model systems.

An important finding was the strong performance of the Den2 5′ UTR, which has been reported as an efficient ribosome recruiter, despite not ranking among the top candidates in massively parallel screening studies.^27^ In this study, Den2 yielded low expression *in vitro* and in zebrafish embryos without a Kozak sequence, yet still outperformed the Syn1 5′ UTR (Figure 2; Figure S2–3). With a Kozak sequence, however, Den2 consistently performed strongly across model systems and coding sequences (Figure 3–5; Figure S3–S6, S9). Interestingly, pairing Den2 with its native Den2 3′ UTR reduced expression relative to combinations with alternative 3′ UTRs, suggesting that the translational performance of this viral-derived 5′ UTR is highly context-dependent. Together, these findings further emphasize that Kozak sequences and UTRs should be considered interacting design features rather than independent optimization parameters in IVT mRNA engineering.

Regarding the synthetic UTRs, Syn2 consistently outperformed Syn1, both *in vitro* and especially in zebrafish embryos. However, in these head-to-head comparisons, we used their original Kozak sequences, which contained an adenosine at position –5 for Syn2, whereas Syn1 carried a guanosine at that position. Notably, substituting A-to-G at –5 in Syn2 significantly reduced expression levels both *in vitro* and in zebrafish embryos (Figure 4; Figure S5–S6), suggesting that the main performance difference stemmed from the nucleotide identity at position –5. Additionally, in mice, the synthetic UTRs were compared with the optimal Kozak sequence and the same HBB 3′ UTR, and they performed comparably (Figure 5; Figure S9), favoring standardized Kozak sequence design in UTR comparisons, and further supporting the benefit of G-to-A substitution at the –5 position.

Notably, several UTR modifications resulted in unanticipated lower expression, even when the optimal Kozak sequence was employed for Syn2 and Den2 (Figure 4; Figure S5–6). This might be partially explained by altered secondary structures upon modification, which appear to reduce accessibility of translation initiation sites (TISs), as suggested by RNAfold structure predictions (Figure S10). When the Den2 5′ UTR is further truncated to 25 nt, these structures appear the least complex, suggesting greater accessibility. Nevertheless, the full-length Den2 5′ UTR, which contains stem–loop structures independent of coding sequence or paired 3′ UTR (Figures S10), performed consistently well, suggesting strong compatibility with eukaryotic translation initiation factors.

Regarding synthetic UTRs, truncation may not be beneficial after rigorous optimization that accounts for uORFs and other undesirable sequence features. Indeed, none of the 5′ UTRs tested here contain uORFs. Truncation of Syn2 indeed appeared to increase the complexity of 5′ UTR secondary structure (Figure S10A). Additionally, in mice, an unanticipated low performer was 25 nt-Den2 with AES-mtRNR1, and to a lesser extent, half-length Den2 with HBB (Figure 5; Figure S10C). Secondary structure, which appeared to reduce accessibility of TISs, might be responsible for these effects (Figure S10C). Although these structure predictions provide some indication of unfavorable structures, it is important to note that they were performed assuming unmodified nucleotides, since no function yet exists to implement 5-moU modifications in mRNA structure prediction.

Altogether, we identify an optimal Kozak sequence that deviates from the consensus at the −5 position. These findings underscore the importance of standardized Kozak sequence design when comparing UTR performance to avoid confounding effects. In addition, our results highlight the need to consider the interplay among UTRs, the coding sequence, and resulting secondary structure in IVT mRNA design. In line with this, recent studies have begun to develop predictive and generative frameworks that integrate these parameters,^58–60^ which are expected to further advance the rational design of potent mRNA therapeutics.

## Materials & Methods

### Reagents

All primers for PCR were obtained from Integrated DNA Technologies B.V. (Leuven, Belgium). New England Biolabs (NEB) reagents for *in vitro* transcription (IVT) and DNA or RNA purification were purchased from Cell Signaling Technology Europe B.V. (Leiden, The Netherlands). TriLink CleanCap® 5-methoxyuridine (5moU) modified EGFP-mRNA, 5-Methoxyuridine-5’-Triphosphate, and CleanCap® AG Reagent were purchased from Tebubio (Heerhugowaard, The Netherlands). Quant-IT™ RiboGreen RNA Assay Kit, 0.5 M Citrate Buffer (pH 4.0), Invitrogen reagents for IVT reactions, IVT mRNA elution, and agarose gel staining, Lipofectamine™ MessengerMAX™ Transfection Reagent, and Gibco supplements for culture media were purchased from Fisher Emergo B.V. (Landsmeer, The Netherlands). Dlin-MC3-DMA and SM-102 were purchased from BroadPharm (San Diego, CA, USA). 1,2-distearoyl-sn-glycerol-3-phosphocholine (DSPC), and 1,2-dimyristoyl-rac-glycero-3-methoxypolyethylene glycol-2000 (DMG-PEG2K) were purchased from Avanti Polar Lipids (Alabaster, USA). Cholesterol (14606-100G-F) was purchased from Merck KGaA (Darmstadt, Germany). The lipid-dye conjugate DiD (1,1’-Dioctadecyl-3,3,3’,3’-Tetramethylindodicarbocyanine, 4-Chlorobenzenesulfonate Salt) was purchased from Fisher Emergo B.V. (Landsmeer, The Netherlands). Tri-GalNAc-PEG2000-DSPE was obtained from Sussex Research Laboratories (Ottawa, ON, Canada). DPBS (10×) was obtained from VWR International B.V. (Amsterdam, The Netherlands). Vivaspin 500 centrifugal filters (MWCO 100K or 300K) were obtained from Sartorius (Göttingen, Germany). RPMI and DMEM culture media and fetal bovine serum were obtained from Capricorn Scientific GmbH (Ebsdorfgrund, Germany). PromoCell Endothelial Cell Growth Medium MV2, WST-1 cell viability reagent, and ampicillin (sodium salt; A8351) were purchased from Merck KGaA (Darmstadt, Germany). Thermo Scientific™ Pierce™ D-Luciferin, Monopotassium Salt was purchased from Life Technologies Corporation (Bleiswijk, The Netherlands). PreciControl ClinChem Multi 2 (05947774) and enzymes (228180; C8868; P8375) for serum cholesterol determination were purchased from Merck KGaA (Darmstadt, Germany).

### IVT mRNA design and template generation

Flanked gene fragments containing a *Mus musculus* Genewiz codon-optimized^27^ red-shifted firefly luciferase coding sequence (amino acid sequence derived from a previous report^50^), the native murine Ldlr-mRNA sequence extracted from the NCBI database (AF425607.1), which was subsequently *Mus musculus* Genewiz codon-optimized, and pUC57-mini vectors containing a T7 promoter with an AGG initiator, human β-globin UTRs or Comirnaty (BNT) UTRs or Den2 UTRs, and a poly(A_35_) sequence inserted at the EcoRV site were obtained from GenScript Biotech B.V. (Rijswijk, The Netherlands). Flanked gBlock DNA fragments of the synthetic UTRs Syn1 (NeoUTR3^26^) and Syn2 (UTR4^19,25^), each with a 5′ part of the EGFP coding sequence, were obtained from Integrated DNA Technologies (Coralville, IA, USA). RNA sequences of the UTRs and coding sequences used in this study are listed in Table S1 and Table S2, respectively. Gene fragments and constructs cloned into the standard vector pUC57-mini are listed in Table S3.

To generate all UTR and CDS combinations, pUC57-mini vectors were linearized by PCR with primers that introduce 15–25 nt overlaps with the insert DNA fragments (if needed), using Q5^®^ High-Fidelity Hot Start DNA polymerase (New England Biolabs, NEB), and purified with the Monarch^®^ PCR and DNA Cleanup kit (NEB). Subsequently, the desired (flanked) DNA fragment or PCR product was cloned into the linearized vector using NEBuilder^®^ HiFi DNA assembly master mix (NEB) and transformed into Stable Competent *E. coli* (NEB). A PCR-prescreened clone was cultured overnight in LB supplemented with ampicillin (100 µg/mL) to amplify the desired plasmid. The plasmid was isolated using the Monarch^®^ Plasmid Miniprep kit (NEB). Subsequently, plasmid sequences were verified by Sanger sequencing performed by BaseClear (Leiden, The Netherlands). Kozak variants were generated by site-directed mutagenesis PCR with primers containing the desired Kozak sequence (as a mutation or tail), using the original plasmids as templates. Correct-sized PCR products were circularized by KLD enzyme mix reactions (NEB), transformed to Stable Competent *E. coli,* amplified, purified, and verified as described above. Sanger sequencing results of the 5′ UTR variants are shown in Figure S12.

Template for *in vitro* transcription (IVT) was generated by PCR using Q5^®^ High-Fidelity Hot Start DNA polymerase (NEB) in a 100 µL reaction using a reverse primer annealing at the 3′ end of the 3′ UTR, installing a poly(A_120_) tail. Then, the size and purity of the PCR-produced IVT template were verified by a 1.2% agarose gel stained with SYBR^TM^ Safe (Invitrogen). Subsequently, any circular plasmid DNA used as the template in the PCR reaction was digested by adding 2 µL of DpnI (NEB) and incubated for 20 minutes at 37°C in a thermal cycler. Finally, the IVT template was purified using the Monarch^®^ PCR and DNA Cleanup kit (NEB), and its concentration was measured with a NanoDrop One (Thermo Fisher Scientific). Primers used in this study are listed in Table S3.

### In vitro transcription and mRNA purification

mRNAs were synthesized by *in vitro* transcription using the following reaction conditions: 25–40 ng/µL purified IVT template, 3.75 U/µL Hi-T7 RNA polymerase (NEB) accompanied with 1× of its supplied buffer, 5 mM ribonucleotides (NEB) in which UTP was replaced with 5moU (TriLink BioTechnologies), 1 U/µL Murine RNase inhibitor (NEB), 4 U/mL Yeast inorganic pyrophosphatase (NEB), 4 mM CleanCap^®^ AG Reagent (TriLink BioTechnologies), 26 mM MgCl_2_, and 5 mM DTT (both Invitrogen). The reaction was incubated for 3 hours at 50°C in a thermal cycler. Afterward, 1 µL of DNase I-XT (NEB) per µg of template DNA was added to the reaction, which was then incubated for 15 min at 37 °C in a thermal cycler. Immediately thereafter, the IVT reaction was purified using the Monarch^®^ Spin RNA Cleanup kit (50 or 500 µg; NEB) and eluted into 1 mM citrate buffer, pH 6.4 (Invitrogen), and kept on ice. The integrity and purity of 100 ng of purified IVT mRNA were verified using a 2% agarose gel stained with SYBR^TM^ Gold (Invitrogen) and the High Range ssRNA ladder (NEB), both diluted in the supplied 2× RNA loading dye, following the manufacturer’s sample preparation recommendations. The IVT mRNA was diluted to 1 mg/ml, snap-frozen on dry ice, and stored at –80 °C until use.

### Cell culture, lipofection, and flow cytometry

The murine immortalized dendritic cell line DC2.4 was kindly provided by Kenneth Rock, University of Massachusetts Medical School, Worcester, MA, USA. The human cell lines HEK293T, HeLa, and Jurkat were obtained from ATCC^®^ (VA, USA). HUVEC cells (C-12203) were purchased from Merck KGaA (Darmstadt, Germany). DC2.4 cells were cultured in RPMI 1640 without L-Glutamine. HEK293T and HeLa were cultured in DMEM High Glucose (4.5 g/L) without L-Glutamine. Both media were supplemented with 10% Advanced heat-inactivated FBS, 1% penicillin-streptomycin, and 2 mM GlutaMAX™. HUVEC cells were cultured in Endothelial Cell Growth Medium MV 2 (PromoCell) supplemented with the supplied supplement mix. Cells were maintained at 37 °C in a humidified atmosphere with 5% CO_2_.

Cells were seeded at 1–2 × 10^4^ cells/well in a 96-well flat-bottom plate and allowed to adhere for 24 hours. Lipofectamine™ MessengerMAX™ reagent diluted in Opti-MEM™ (Gibco) without supplements was mixed with 600–900 ng mRNA diluted in Opti-MEM™ at the manufacturer’s recommended ratio. The mixtures were directly vortexed for 60 seconds to generate lipoplexes. The original medium of cells seeded in flat-bottom 96-well plates was replaced with Opti-MEM™ supplemented with 2.5% Advanced heat-inactivated FBS and 1% penicillin-streptomycin. Cells were then treated with lipoplexes at mRNA concentrations of 100, 250, and 500 ng mL^-1^ in supplemented medium and incubated for 24 hours at 37 °C in a 5% CO_2_ atmosphere. At least two different cell lines were transfected with the same lipoplexes per experiment.

After incubation, cells were washed with PBS, resuspended in FACS buffer (PBS, 2% FBS, 0.1% sodium azide, and 1 mM EDTA), and analyzed by flow cytometry (CytoFLEX S, Beckman Coulter, CA, USA) for EGFP fluorescence. Data were analyzed using FlowJo software V10 (Tree Star Inc., OR, USA). The geometric mean fluorescence intensity (gMFI) of EGFP was graphed using GraphPad Prism 10.

### Lipid nanoparticle manufacturing

Lipid nanoparticles were self-assembled as previously described^61^ with minor adaptations. Lipids were combined at the desired molar ratios and concentrations from stock solutions (0.05–10 mM) in chloroform. Lipid molar ratios and characterization data for the LNP formulations used in this study are listed in Table S4. Briefly, for zebrafish embryo experiments, an Onpattro-like formulation containing Dlin-MC3-DMA and 10% DSPC was used, whereas for the mice experiments, a formulation containing SM-102, 10% DSPC, and 0.05% DSPE-PEG2K-triGalNAc was employed. Solvents were evaporated under a nitrogen flow. The lipid film was dissolved in absolute ethanol to a total lipid concentration of 4–6 mM. 1 mg/mL RNA stored at –80 °C was thawed and diluted in 25 mM citrate buffer to a final mRNA concentration of 33.33–50 µg/mL. Depending on the amount of mRNA to be encapsulated, volumes ranged from 100–300 µL and 300–900 µL for the lipid and mRNA solutions, respectively. The solutions were loaded into two separate sterile syringes, connected to a T-junction microfluidic mixer, and mixed using two Chemyx Fusion 100X syringe pumps at a 3:1 (RNA:lipid) flow ratio, yielding a total flow rate of 2.4 mL/min. After mixing, the solution was loaded directly into a dialysis cassette (Slide-A-Lyzer™ 20 kDa MWCO, 0.5 mL, Thermo Scientific) and dialyzed against PBS for 2–16 hours. LNPs were concentrated using Vivaspin 500 centrifugal filters at 2000 × g at 4 °C.

### Lipid nanoparticle characterization

LNP sizes and zeta potentials were measured using a Malvern Zetasizer Nano ZS (software version 7.13, Malvern Panalytical). For DLS (operating wavelength = 633 nm), measurements were performed at 25 °C in 1x PBS (pH = 7.4) at a total lipid concentration of approximately 0.1 mM. Zeta potentials were measured at 0.5 mM total lipid concentration, using a folded capillary cell (DTS1070, Malvern), at room temperature. All reported DLS measurements and zeta potentials are the average of three measurements.

Encapsulation efficiency and encapsulated RNA concentration were analyzed using the Quant-iT^TM^ Ribogreen RNA assay by diluting 1–2 µL of concentrated LNPs back to their original concentration after dialysis in 1× PBS, and then following the manufacturer’s low-range assay protocol. RNA encapsulation efficiency was calculated using the following equation:

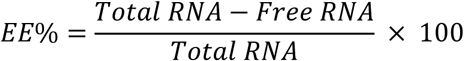

### Animal experiments and husbandry

All animal work was approved by the Leiden University Animal Ethics Committee, and the animal experiments were performed in accordance with the guidelines from Directive 2010/63/EU of the European Parliament on the protection of animals used for scientific purposes. *Ldlr ^-/-^* mice were purchased from Jackson Laboratory (CA, USA), bred in-house, kept under standard laboratory conditions, and provided with food (chow) and water *ad libitum*.

Zebrafish were maintained in accordance with standard zebrafish rearing protocols (https://zfin.org), which comply with the EU Animal Protection Directive 2010/63/EU. Housing and husbandry recommendations were followed as previously recommended.^62^ Fertilization was performed by natural spawning at the beginning of the light period, and eggs were raised at 28°C in egg water (60 μg/ml Instant Ocean sea salts). Only wild-type AB/TL zebrafish were used in this study.

### Zebrafish microinjections, confocal imaging, and image processing

LNP formulations were injected into zebrafish embryos at time points ranging from 54–60 hours post-fertilization (hpf) using a modified microangraphy protocol.^63^ One nanoliter volume of nanoparticle formulation was calibrated and injected into the duct of Cuvier after the embryos were embedded in 0.4% agarose containing 0.01% tricaine as described.^39^

At least three randomly selected, well-injected embryos were mounted in a 60 mm confocal dish with 0.4% agarose in egg water containing 0.01% buffered 3-aminobenzoic acid ethyl ester (tricaine; Sigma-Aldrich, A-5040) and imaged at 24 hours post-injection (hpi). Three to six whole embryos were imaged per group.

Confocal z-stacks were captured using a Leica TCS SP8 or a Leica STELLARIS 5 confocal laser-scanning microscope, equipped with a 10× air objective (HCX PL FLUOTAR) for whole-body imaging. To ensure comparability across experiments, microscopy settings, including laser intensity, gain, and offset, were kept constant across stacks and sessions. Additionally, when groups were compared across experiments (such as in Figure 4), the same microscope with identical settings was used. Whole-embryo images consisted of 3–4 overlapping z-stacks were stitched using FIJI imaging software.^64^

Whole-embryo EGFP CTCF was measured using FIJI (ImageJ) by selecting the sum of slices’ z-projection of the entire embryo with the wand tool, then measuring MFI, area, and integrated density of the selected region, along with background fluorescence MFI outside the embryo. CTCF was calculated, and background fluorescence was subtracted by computing (integrated density) – (area × background MFI).

### In vivo luciferase expression and IVIS imaging

16-week-old male low-density lipoprotein-deficient (*Ldlr ^-/-^*) mice (n = 4 per group) received 0.15 mg/kg of FLuc-mRNA encapsulated in triGalNAc-LNPs (Table S4) intravenously via tail vein injection in a 100 µL volume in PBS. Six hours later, they received a second 100 µL tail vein injection of a 4.5 mg/mL D-luciferin solution in Dulbecco’s PBS (150 mg/kg) and were allowed to recover for 3 minutes. Subsequently, mice were anesthetized with isoflurane in oxygen (induction 3%, maintenance 2%) and imaged in a light-tight IVIS system on a warmed stage at 5 minutes post-D-luciferin injection, with a constant 5-second exposure time. For a subset of mice, the D-luciferin injection and imaging procedure was repeated at 24 h post LNP administration.

Live bioluminescence images were analyzed using the PerkinElmer Living Image software (v4.7.3). The radiance (photons/sec/cm^2^/sr) scale was adjusted across all images to the range 5×10^6^–5×10^7^ for comparison. ROIs were selected using a fixed threshold of 10 or manually adjusted to the organ size, since the resulting ROI size was often not representative of the organ. ROIs at the injection site were ignored if below 5×10^6^ or added to the other ROI value for that individual. Overall data trends were the same for both ROI selection methods; therefore, manual ROI selection and threshold setting were employed, as they better reflected organ size and intergroup differences. ROI values were measured and exported, and total flux per second was plotted using GraphPad Prism 10.

### Consensus and optimized Kozak sequence comparison

Natural vertebrate Kozak sequences were derived from annotated coding transcripts from human, mouse, and zebrafish transcriptomes. For each transcript, the nucleotide window spanning positions −9 to −1 relative to the annotated start codon (AUG) was extracted from the corresponding cDNA sequence to generate species-specific Kozak datasets, which were subsequently combined into a vertebrate reference set. Experimentally tested Kozak variants were ranked by relative protein expression (Table S5), and sequences were represented in weighted FASTA files with linear scaling of normalized expression values. Nucleotide frequencies at each position were calculated from the weighted Kozak dataset and compared to frequencies in the vertebrate reference dataset. Enrichment was calculated as:

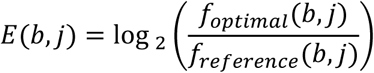

where b denotes the nucleotide and j the position relative to the start codon. If a nucleotide was absent from the experimentally derived Kozak dataset at a given position f_optimal_(*b, j*) = 0, the enrichment value approaches −∞; for visualization purposes, enrichment values were capped at ±3. Sequence logos were generated using WebLogo version 3.9.0; https://weblogo.threeplusone.com/create.cgi). Heatmaps were generated with GraphPad Prism 10.

### In vivo Ldlr replacement

16-week-old male *Ldlr ^-/-^* mice were fed a Western-type diet starting 14 days before LNP treatment to elevate serum cholesterol levels. Three days before LNP treatment, mice were bled to measure cholesterol levels, allowing normalization to each individual’s baseline throughout the experiment. Mice then received a single 0.25 mg/kg dose of Ldlr-mRNA encapsulated in triGalNAc-LNPs (Table S4) via tail vein injection in a 100 µL volume in PBS. Subsequently, mice were bled on days 1, 4, 7, and 14 after LNP administration to measure serum cholesterol levels.

### Serum cholesterol determination

Serum was obtained by collecting whole blood in cloth-inducing microvette CB300Z tubes (Sarstedt), followed by centrifugation at 2000 × g for 10 minutes at 4 °C. Serum was stored at −80 °C until further usage. Total cholesterol levels in serum were measured by performing an enzymatic colorimetric analysis (Roch/Hitachi, Germany), using Precipath standardized serum (1.69 mg/mL; Roche/Hitachi) as an internal standard.

### Statistical analysis

In case of 3 data points per group, between-group comparisons were performed using Brown-Forsythe & Welch ANOVA with Dunnett T3 correction for multiple comparisons. One-way ANOVA with Tukey’s correction for multiple comparisons was performed when there were more than 3 data points per group, with the note that statistical tests with 4 data points were intended to be exploratory. A two-way ANOVA with Dunnett correction for multiple comparisons was performed exploratorily to compare serum cholesterol levels over time between groups. QQ-plots were computed to investigate whether the data followed a Gaussian distribution. All statistical analyses were performed with GraphPad Prism 10.

## Data availability statement

Data supporting the findings of this study are available from the corresponding authors upon reasonable request.

## Acknowledgements

B.S and R.K. receive funding from ERA4Health and the Dutch Research Council (grant number era4healthcvd-112). A.K. has received funding from the European Research Council (ERC) under the European Union’s Horizon Europe research and innovation programme (grant agreement No. 101118999, Cat4CanCenter, ERC-2023-SyG). Parts of Figures 1, 2, and 5 were created with BioRender.com using an academic lab subscription. R.K. kindly acknowledges Ye Zeng for assistance in preliminary *in vitro* experiments.

## Author contributions

Conceptualization, R.K., T.F., and B.S.; Methodology, R.K., T.F., and B.S.; Investigation, R.K., T.F., E.V., and B.S.; Formal analysis, R.K., T.F., E.V., and B.S.; Data curation, R.K.; Resources, B.S. and A.K.; Visualization, R.K., T.F., and B.S.; Supervision, B.S. and A.K.; Project administration, R.K.; Funding acquisition, A.K. and B.S.; Writing – original draft, R.K.; Writing – review & editing, all authors. B.S. and A.K. share senior authorship. All authors reviewed and approved the final manuscript.

## Declaration of interests

The authors declare no competing interests.

## Notes

### Competing Interest Statement

The authors have declared no competing interest.

