## Supplemental Material for "A single-nucleotide change in the Kozak sequence enhances protein expression"

### Supplemental Information

**Table S1. UTR sequences used in this study.**

| | Name (original name) | RNA sequence (5′ – 3′) | Length (nt) |
|---|---|---|---|
| 5′ UTRs | HBB[1] (human β-globin) | CAGGGCAGAGCCAUCUAUUGCUUACAUUUGCUUCUGACACAACUGUGUUCACUAGCAACCUCA<br>AACAGACACC | 73 |
| | Syn1 (NeoUTR3[2]) | CUUGUCUCGCUCCGGGGAACGCUCGGAAACUCCCGGCCGCCGCCACCCGCGUCUGUUCUGUU<br>ACACAAGGGAAGAAAAGCCGCUGCCGCACUCCGAGUGUGCCGCCACC | 100 (+ 9) |
| | Syn2 (UTR4[3,4]) | CUGAAACACGGUGGAGAGUUUAUUGCAAAAUAACGCGUCCAUUCGACACC | 50 |
| | Den2[5] | AGUUGUUAGUCUACGUGGACCGACAAAGACAGAUUCUUUGAGGGAGCUAAGCUCAACGUAGUU<br>CUAACAGUUUUUUAAUUAGAGAGCAGAUCUCUG | 96 |
| | HBA[6] | GAAUAAACUAGUAUUCUUCUGGUCCCCACAGACUCAGAGAGAACCCGCCACC | 52 |
| 3′ UTRs | HBB[1] (human β-globin) | GCUCGCUUUCUUGCUGUCCAAUUUCUAUUAAAGGUUCCUUUGUUCCCUAAGUCCAACUACUAA<br>ACUGGGGGAUAUUAUGAAGGGCCUUGAGCAUCUGGAUUCUGCCUAAUAAAAAACAUUUAUUUU<br>CAUUGC | 132 |
| | mtRNR1[7] | CAAGCACGCAGCAAUGCAGCUCAAAACGCUUAGCCUAGCCACACCCCCACGGGAAACAGCAGU<br>GAUUAACCUUUAGCAAUAAACGAAAGUUUAACUAAGCUAUACUAACCCCAGGGUUGGUCAAUU<br>UCGUGCCAGCCACACC | 142 |
| | AES-mtRNR1[6,7] | CUGGUACUGCAUGCACGCAAUGCUAGCUGCCCCUUUCCCGUCCUGGGUACCCCGAGUCUCCC<br>CCGACCUCGGGUCCCAGGUAUGCUCCCACCUCCACCUGCCCCACUCACCACCUCUGCUAGUU<br>CCAGACACCUCCCAAGCACGCAGCAAUGCAGCUCAAAACGCUUAGCCUAGCCACACCCCCACG<br>GGAAACAGCAGUGAUUAACCUUUAGCAAUAAACGAAAGUUUAACUAAGCUAUACUAACCCCAGG<br>GUUGGUCAAUUUCGUGCCAGCCACACC | 278 |
| | Den2[5] | AAAGCAAAACUAACAUGAAACAAGGCUAGAAGUCAGGUCGGAUUAAGCCAUAGUACGGAAAAAA<br>CUAUGCUACCUGUGAGCCCCGUCCAAGGACGUUAAAAGAAGUCAGGCCAUCAUAAAUGCCAUA<br>GCUUGAGUAAACUAUGCAGCCUGUAGCUCCACCUGAGAAGGUGUAAAAAAUCCGGGAGGCCAC<br>AAACCAUGGAAGCUGUACGCAUGGCGUAGUGGACUAGCGGUUAGAGGAGACCCCUCCCUUAC<br>AAAUCGCAGCAACAAUGGGGGCCCAAGGCGAGAUGAAGCUGUAGUCUCGCUGGAAGGACUAG<br>AGGUUAGAGGAGACCCCCCCGAAACAAAAAACAGCAUAUUGACGCUGGGAAAGACCAGAGAUC<br>CUGCUGUCUCCUCAGCAUCAUUCCAGGCACAGAACGCCAGAAAAUGGAAUGGUGCUGUUGAAU<br>CAACAGGUUCU | 451 |

**Table S2. Coding sequences used in this study.**

| Name | RNA sequence (5′ – 3′) | Length (nt) |
|---|---|---|
| EGFP | AUGGUGAGCAAGGGCGAGGAGCUGUUCACCGGGGUGGUGCCCAUCCUGGUCGAGCUGGACGGCGACGUAAACGG<br>CCACAAGUUCAGCGUGUCCGGCGAGGGCGAGGGCGAUGCCACCUACGGCAAGCUGACCCUGAAGUUCAUCUGCAC<br>CACCGGCAAGCUGCCCGUGCCCUGGCCCACCCUCGUGACCACCCUGACCUAUGGAGUGCAGUGCUUCAGCCGCUA<br>CCCCGACCACAUGAAGCAGCACGACUUCUUCAAGUCCGCCAUGCCCGAAGGCUACGUCCAGGAGCGCACCAUCUUC<br>UUCAAGGACGACGGCAACUACAAGACCCGCGCCGAGGUGAAGUUCGAGGGCGACACCCUGGUGAACCGCAUCGAG<br>CUGAAGGGCAUCGACUUCAAGGAGGACGGCAACAUCCUGGGGCACAAGCUGGAGUACAACUACAACAGCCACAACG<br>UCUAUAUCAUGGCCGACAAGCAGAAGAACGGCAUCAAGGUGAACUUCAAGAUCCGCCACAACAUCGAGGACGGCAG<br>CGUGCAGCUCGCCGACCACUACCAGCAGAACACCCCCAUCGGCGACGGCCCCGUGCUGCUGCCCGACAACCACUA<br>CCUGAGCACCCAGUCCGCCCUGAGCAAAGACCCCAACGAGAAGCGCGAUCACAUGGUCCUGCUGGAGUUCGUGAC<br>CGCCGCCGGGAUCACUCUCGGCAUGGACGAGCUGUACAAGUAA | 720 |
| FLuc | AUGGAGGACGCCAAGAACAUCAAGAAGGGCCCUGCCCCUUUCUACCCUCUGGAGGACGGCACCGCCGGCGAGCAG<br>CUGCACAAAGCCAUGAAGAGGUACGCUCUGGUGCCUGGCACCAUCGCCUUCACCGACGCCCACAUCGAGGUGGAC<br>AUCACCUACGCCGAGUACUUCGAGAUGAGCGUGAGGCUGGCCGAGGCAAUGAAGCGUUAUGGCCUGAACACCAAC<br>CAUAGGAUCGUGGUGUGCAGCGAGAACAGCCUGCAGUUCUUCAUGCCUGUGCUGGGCGCCCUGUUCAUCGGCGU<br>GGCCGUUGCCCCUGCCAACGACAUCUACAACGAGAGGGAGCUGCUGAACAGCAUGGGCAUCUCUCAGCCUACCGU<br>AGUGUUCGUGAGUAAAAAGGGCCUGCAGAAGAUCCUGAACGUGCAGAAGAAGCUGCCUAUCAUUCAGAAGAUCAUC<br>AUCAUGGACAGCAAGACCGACUACCAAGGCUUUCAGAGCAUGUACACCUUCGUGACAAGCCACCUGCCUCCUGGCU<br>UCAACGAGUACGACUUCGUGCCUGAGAGCUUCGAUAGGGACAAGACCAUCGCCCUGAUCAUGAACAGCAGCGGCA<br>GCACCGGCCUGCCUAAGGGCGUGGCCCUGCCUCAUAGGGCCUUAUGCGUGAGGUUUAGUCACGCUAGGGACCCU<br>AUCUUCGGCAAUCAGAUCGCCCCUGACACCGCCAUCCUGAGCGUCGUCCCUUUCCACCACGGCUUCGGCAUGUUC<br>ACCACCCUGGGCUACCUGAUCUGCGGCUUUAGGGUGGUGCUGAUGUAUAGGUUCGAGGAGGAGCUGUUCCUGAG<br>GAGCCUGCAAGACUACAAGAUUCAGACCGCCCUGCUGGUGCCUACCCUGUUCAGCUUCCUGGCCAAGAGCACCCU<br>GAUCGACAAGUAUGAUCUAUCCAACCUGCACGAGAUCGCAAGCGGGGGCGCACCUCUGAGCAAAGAGGUGGGCGA<br>GGCCGUGGCCAAGAGGUUCCACCUGCCUGGCAUUAGGCAAGGCUACGGCCUGACCGAAACCACAAGCGCCAUCCU<br>GAUCACCCCUAAGGGCGAUGAUAAACCUGGCGCCGUGGGCAAGGUGGUACCUUUCUUCGAAGCCAAGGUGGUGGA<br>CCUGGACACCGGCAAGACCCUGGGCGUGAAUCAGAGGGGCGAGCUGUGUGUUAGGGGCCCUAUGAUCAUGAGCG<br>GCUACGUGAACAACCCUGAGGCCACCAACGCCCUCAUCGAUAAGGAUGGAUGGCUGCACAGCGGCGACCUGGCCU<br>ACUGGGACGAGGACGAGCACUUCUUCAUCGUGGGAAGGCUGAAGAGCCUGAUCAAGUACAAGGGCUACCAAGUGG<br>CCCCUGCCGAGCUUGAGAGCAUCCUGCUGCAGCACCCUAACAUCUUUGAUGCCGGCGUUGCGGGGUUGCCGGAU<br>GACGACGCCGGCGAACUCCCUGCCGCCGUGGUGGUGCUGGAGCACGGCAAGACCAUGACCGAGAAGGAGAUCGU<br>GGACUACGUGGCUAGCCAAGUGACCACCGCCAAGAAGCUGAGGGGCGGCGUGGUUUUUGUUGACGAAGUGCCUAA<br>GGGCCUGACCGGCAAGAGGGACGCUAGGAAGAUUAGGGAGAUCCUGAUCAAGGCCAAGAAGGGCGGCAAGAUCGC<br>CGUGUGA | 1653 |
| Ldlr | AUGAGCACCGCCGACCUGAUGCGAAGGUGGGUGAUCGCCUUGCUACUGGCUGCCGCCGGAGUGGCCGCCGAGGA<br>CAGCUGCUCUAGGAACGAAUUUCAGUGUAGGGACGGCAAGUGCAUCGCUAGCAAGUGGGUCUGCGACGGCUCACC<br>UGAAUGCCCUGACGGCAGCGACGAGAGCCCUGAAACCUGCAUGAGCGUGACCUGUCAGAGCAAUCAGUUCAGCUG<br>CGGCGGAAGGGUGUCUAGGUGCAUCCCUGACAGCUGGAGGUGCGACGGCCAAGUGGACUGCGAGAACGACAGCG<br>ACGAGCAAGGCUGCCCUCCUAAGACCUGCUCACAAGACGACUUUAGGUGCCAAGACGGGAAAUGUAUCAGUCCUCA<br>GUUUGUUUGUGACGGUGAUCGGGACUGUUUAGACGGCAGCGAUGAAGCCCACUGUCAAGCAACCACCUGCGGCCC<br>UGCCCACUUUAGGUGCAACAGCAGCAUCUGCAUCCCUAGCCUGUGGGCCUGCGAUGGCGACGUGGACUGCGUGGA<br>UGGAAGCGAUGAGUGGCCUCAGAACUGCCAAGGAAGGGACACCGCUAGCAAGGGCGUGAGCAGCCCUUGCAGCAG<br>CCUGGAGUUCCACUGCGGCAGCAGCGAGUGCAUCCAUAGGAGCUGGGUCUGUGAUGGUGAGGCGGAUUGUAAGG<br>ACAAGAGCGACGAAGAACAUUGCGCCGUGGCCACCUGUAGGCCUGACGAAUUUCAGUGCGCAGACGGCAGCUGCA<br>UCCACGGCUCUAGGCAGUGCGAUAGGGAGCACGACUGUAAGGAUAUGAGUGACGAGCUGGGCUGCGUGAACGUGA<br>CACAGUGCGACGGCCCUAACAAGUUCAAGUGCCACAGCGGCGAAUGCAUCAGCCUGGACAAGGUGUGCGACAGCG<br>CUAGGGACUGCCAAGACUGGAGCGACGAGCCUAUCAAGGAGUGCAAGACCAACGAGUGCCUGGACAACAACGGCG<br>GCUGCAGCCACAUCUGCAAGGACCUGAAGAUCGGCAGCGAGUGCCUGUGCCCUAGCGGCUUUAGGCUGGUGGACC<br>UGCAUAGGUGCGAGGACAUCGACGAGUGCCAAGAGCCUGACACCUGCUCUCAGCUGUGCGUGAACCUGGAGGGCA<br>GCUACAAGUGCGAGUGCCAAGCCGGCUUCCACAUGGACCCUCACACAAGGGUGUGCAAGGCCGUGGGCAGCAUCG<br>GCUACCUGCUGUUCACCAAUAGGCACGAGGUGAGGAAGAUGACCCUGGAUAGGAGCGAGUACACAAGCUUACUGC<br>CUAAUCUGAAGAACGUGGUGGCCCUGGACACCGAGGUGACCAACAAUAGGAUCUACUGGAGCGACCUGUCUCAGA<br>AGAAGAUCUACAGCGCCCUGAUGGACCAAGCCCCUAACCUGAGCUACGACACCAUCAUCUCUGAAGACCUGCACGC<br>CCCUGACGGCCUGGCCGUGGACUGGAUCCAUAGGAACAUCUACUGGACUGACAGCGUGCCUGGCAGCGUGAGCGU<br>GGCCGACACCAAGGGCGUGAAGAGGAGGACCCUGUUCCAAGAGGCCGGCUCUAGGCCUAGGGCCAUCGUGGUGG<br>ACCCUGUGCACGGCUUCAUGUACUGGACCGAUUGGGGCACCCCUGCCAAGAUCAAGAAGGGCGGCCUGAACGGCG<br>UGGACAUCCACAGCCUGGUGACCGAGAACAUUCAGUGGCCUAACGGCAUCACCCUGGACCUGAGCAGCGGAAGGC<br>UGUACUGGGUGGACAGCAAGCUGCACAGCAUCAGCAGCAUCGACGUGAACGGCGGCAAUAGGAAGACCAUCCUGG<br>AGGACGAAAAUAGGCUGGCCCACCCUUUCAGCCUGGCCAUCUACGAGGACAAGGUGUACUGGACUGAUGUGAUUA<br>ACGAGGCCAUCUUCAGCGCCAAUAGGCUGACCGGCAGCGACGUGAACCUGGUGGCCGAGAACCUGCUGAGCCCUG<br>AGGACAUCGUGCUGUUCCACAAGGUGACACAGCCUAGGGGCGUGAACUGGUGCGAAACCACCGCCCUUCUGCCUA<br>ACAGCGGCUGUCAGUACCUGUGCCUGCCUGCCCCUCAGAUCGGCCCUCACAGCCCUAAGUUCACCUGCGCCUGCC<br>CUGAUGGCAUGCUACUCGCCGAGGACAUGAGGAGCUGCCUGACGGAGGUGGACACCGUGCUGACCACUCAAGGCA<br>CCUCAGCAGUGAGGCCUGUGGUGACCGCUAGCGCCACAAGGCCUCCUAAGCACUCUGAGGACCUCAGCGCCCCUA<br>GCACCCCGAGGCAGCCUGUGGACACCCCUGGCCUGAGCACCGUGGCCUCUGUGACCGUGAGCCACCAAGUGCAAG<br>GCGACAUGGCGGGGAGGGGCAAUGAGGAGCAGCCGCACGGCGUGAGGUUCCUGAGCAUCUUCUUCCCUAUCGCC<br>CUGGUGGCCCUGCUGGUGCUGGGCGCCGUGCUGCUGUGGAGGAACUGGAGGCUGAAGAACAUCAACAGCAUCAA<br>CUUCGACAACCCUGUGUAUCAGAAGACCACCGAGGACGAGCUGCACAUCUGUAGGAGCCAAGACGGCUACACCUAC<br>CCUUCUAGGCAGAUGGUGAGCCUCGAAGACGACGUAGCCUGA | 2589 |

**Table S3. T7AGG-UTR-pA constructs, (flanked) DNA fragments, and primers used in this study.** Grey nucleotides represent non-binding primer tails.

| Name | Sequence (5′ – 3′) |
|---|---|
| T7AGG-HBB-HBB-pA<br>(cloned into pUC57-mini) | CCAATGATGCTCTAATACGACTCACTATAAGGCAGGGCAGAGCCATCTATTGCTTACATTTGCTTC<br>TGACACAACTGTGTTCACTAGCAACCTCAAACAGACACCTTAAGCTCGCTTTCTTGCTGTCCAATT<br>TCTATTAAAGGTTCCTTTGTTCCCTAAGTCCAACTACTAAACTGGGGGATATTATGAAGGGCCTTG<br>AGCATCTGGATTCTGCCTAATAAAAAACATTTATTTTCATTGCAAAAAAAAAAAAAAAAAAAAAAAAA<br>AAAAAAAAAATCTAGATGATCATCGCCATGTACAGGTACCGCTAGCTATGGACCTGAT |
| T7AGG-HBA-AES-<br>mtRNR1-pA<br>(cloned into pUC57-mini) | CCAATGATGCTCTAATACGACTCACTATAAGGGAATAAACTAGTATTCTTCTGGTCCCCACAGACTC<br>AGAGAGAACCCGCCACCTTAAGCTGGTACTGCATGCACGCAATGCTAGCTGCCCCTTTCCCGTC<br>CTGGGTACCCCGAGTCTCCCCCGACCTCGGGTCCCAGGTATGCTCCCACCTCCACCTGCCCCA<br>CTCACCACCTCTGCTAGTTCCAGACACCTCCCAAGCACGCAGCAATGCAGCTCAAAACGCTTAG<br>CCTAGCCACACCCCCACGGGAAACAGCAGTGATTAACCTTTAGCAATAAACGAAAGTTTAACTAA<br>GCTATACTAACCCCAGGGTTGGTCAATTTCGTGCCAGCCACACCCTGGAAAAAAAAAAAAAAAAA<br>AAAAAAAAAAAAAAAAAATCTAGATGATCATCGCCATGTACAGGTACCGCTAGCTATGGACCTGAT |
| T7AGG-Den2-Den2-pA<br>(cloned into pUC57-mini) | CCAATGATGCTCTAATACGACTCACTATAAGGAGTTGTTAGTCTACGTGGACCGACAAAGACAGAT<br>TCTTTGAGGGAGCTAAGCTCAACGTAGTTCTAACAGTTTTTTAATTAGAGAGCAGATCTCTGATGG<br>TGAGCAAGGGCGAGGAGCATGGACGAGCTGTACAAGTAAAAAGCAAAACTAACATGAAACAAGG<br>CTAGAAGTCAGGTCGGATTAAGCCATAGTACGGAAAAAACTATGCTACCTGTGAGCCCCGTCCAA<br>GGACGTTAAAAGAAGTCAGGCCATCATAAATGCCATAGCTTGAGTAAACTATGCAGCCTGTAGCTC<br>CACCTGAGAAGGTGTAAAAAATCCGGGAGGCCACAAACCATGGAAGCTGTACGCATGGCGTAGT<br>GGACTAGCGGTTAGAGGAGACCCCTCCCTTACAAATCGCAGCAACAATGGGGGCCCAAGGCGA<br>GATGAAGCTGTAGTCTCGCTGGAAGGACTAGAGGTTAGAGGAGACCCCCCCGAAACAAAAAACA<br>GCATATTGACGCTGGGAAAGACCAGAGATCCTGCTGTCTCCTCAGCATCATTCCAGGCACAGAA<br>CGCCAGAAAATGGAATGGTGCTGTTGAATCAACAGGTTCTAGAAAAAAAAAAAAAAAAAAAAAAA<br>AAAAAAAAAAAAATGGTCTTCGTCATCGCCATGTACAGGTACCGCTAGCTATGGACCTGAT |
| Syn1_5′EGFP | gctctaatacgactcactataaggCTTGTCTCGCTCCGGGGAACGCTCGGAAACTCCCGGCCGCCGCCAC<br>CCGCGTCTGTTCTGTTACACAAGGGAAGAAAAGCCGCTGCCGCACTCCGAGTGTGCCGCCACC<br>atggtgagcaagggcgaggagctgttcaccggggtggtgcccatcctggtcgagctggacggcgacgtaaacggccacaagttcagcgtg<br>tccggcgagggcgagggcgatgccacctacggcaagctgaccctgaagttcatctgcaccaccggcaagctgcccgtgccctggcccacc<br>ctcgtgaccaccctgacctatggagtgcagtgcttcagccgctaccccgaccacatgaagcagcacgacttcttcaagtccgccatgcccga<br>aggctacgtccaggagcgcaccatcttcttcaaggacgacg |
| Syn2_5′EGFP | gctctaatacgactcactataaggCTGAAACACGGTGGAGAGTTTATTGCAAAATAACGCGTCCATTCGACAC<br>Catggtgagcaagggcgaggagctgttcaccggggtggtgcccatcctggtcgagctggacggcgacgtaaacggccacaagttcagcgt<br>gtccggcgagggcgagggcgatgccacctacggcaagctgaccctgaagttcatctgcaccaccggcaagctgcccgtgccctggcccac<br>cctcgtgaccaccctgacctatggagtgcagtgcttcagccgctaccccgaccacatgaagcagcacgacttcttcaagtccgccatgcccg<br>aaggctacgtccaggagcgcaccatcttcttcaaggacgacg |
| MmFLuc | aattagagagcagatctctggccgacaccATGGAGGACGCCAAGAACATCAAGAAGGGCCCTGCCCCTTTCTACCCTCTG<br>GAGGACGGCACCGCCGGCGAGCAGCTGCACAAAGCCATGAAGAGGTACGCTCTGGTGCCTGGCACCATCG<br>CCTTCACCGACGCCCACATCGAGGTGGACATCACCTACGCCGAGTACTTCGAGATGAGCGTGAGGCTGGCC<br>GAGGCAATGAAGCGTTATGGCCTGAACACCAACCATAGGATCGTGGTGTGCAGCGAGAACAGCCTGCAGTTC<br>TTCATGCCTGTGCTGGGCGCCCTGTTCATCGGCGTGGCCGTTGCCCCTGCCAACGACATCTACAACGAGAG<br>GGAGCTGCTGAACAGCATGGGCATCTCTCAGCCTACCGTAGTGTTCGTGAGTAAAAAGGGCCTGCAGAAGAT<br>CCTGAACGTGCAGAAGAAGCTGCCTATCATTCAGAAGATCATCATCATGGACAGCAAGACCGACTACCAAGGC<br>TTTCAGAGCATGTACACCTTCGTGACAAGCCACCTGCCTCCTGGCTTCAACGAGTACGACTTCGTGCCTGAG<br>AGCTTCGATAGGGACAAGACCATCGCCCTGATCATGAACAGCAGCGGCAGCACCGGCCTGCCTAAGGGCGT<br>GGCCCTGCCTCATAGGGCCTTATGCGTGAGGTTTAGTCACGCTAGGGACCCTATCTTCGGCAATCAGATCGCC<br>CCTGACACCGCCATCCTGAGCGTCGTCCCTTTCCACCACGGCTTCGGCATGTTCACCACCCTGGGCTACCTG<br>ATCTGCGGCTTTAGGGTGGTGCTGATGTATAGGTTCGAGGAGGAGCTGTTCCTGAGGAGCCTGCAAGACTAC<br>AAGATTCAGACCGCCCTGCTGGTGCCTACCCTGTTCAGCTTCCTGGCCAAGAGCACCCTGATCGACAAGTAT<br>GATCTATCCAACCTGCACGAGATCGCAAGCGGGGGCGCACCTCTGAGCAAAGAGGTGGGCGAGGCCGTGG<br>CCAAGAGGTTCCACCTGCCTGGCATTAGGCAAGGCTACGGCCTGACCGAAACCACAAGCGCCATCCTGATCA<br>CCCCTAAGGGCGATGATAAACCTGGCGCCGTGGGCAAGGTGGTACCTTTCTTCGAAGCCAAGGTGGTGGAC<br>CTGGACACCGGCAAGACCCTGGGCGTGAATCAGAGGGGCGAGCTGTGTGTTAGGGGCCCTATGATCATGAG<br>CGGCTACGTGAACAACCCTGAGGCCACCAACGCCCTCATCGATAAGGATGGATGGCTGCACAGCGGCGACC<br>TGGCCTACTGGGACGAGGACGAGCACTTCTTCATCGTGGGAAGGCTGAAGAGCCTGATCAAGTACAAGGGC<br>TACCAAGTGGCCCCTGCCGAGCTTGAGAGCATCCTGCTGCAGCACCCTAACATCTTTGATGCCGGCGTTGCG<br>GGGTTGCCGGATGACGACGCCGGCGAACTCCCTGCCGCCGTGGTGGTGCTGGAGCACGGCAAGACCATGA<br>CCGAGAAGGAGATCGTGGACTACGTGGCTAGCCAAGTGACCACCGCCAAGAAGCTGAGGGGCGGCGTGGT<br>TTTTGTTGACGAAGTGCCTAAGGGCCTGACCGGCAAGAGGGACGCTAGGAAGATTAGGGAGATCCTGATCAA<br>GGCCAAGAAGGGCGGCAAGATCGCCGTGTGAgctcgctttcttgctgtcc |

| | |
|---|---|
| MmLDLR | aattagagagcagatctctggccgacaccATGAGCACCGCCGACCTGATGCGAAGGTGGGTGATCGCCTTGCTACTGGCT<br>GCCGCCGGAGTGGCCGCCGAGGACAGCTGCTCTAGGAACGAATTTCAGTGTAGGGACGGCAAGTGCATCGC<br>TAGCAAGTGGGTCTGCGACGGCTCACCTGAATGCCCTGACGGCAGCGACGAGAGCCCTGAAACCTGCATGA<br>GCGTGACCTGTCAGAGCAATCAGTTCAGCTGCGGCGGAAGGGTGTCTAGGTGCATCCCTGACAGCTGGAGG<br>TGCGACGGCCAAGTGGACTGCGAGAACGACAGCGACGAGCAAGGCTGCCCTCCTAAGACCTGCTCACAAGA<br>CGACTTTAGGTGCCAAGACGGGAAATGTATCAGTCCTCAGTTTGTTTGTGACGGTGATCGGGACTGTTTAGAC<br>GGCAGCGATGAAGCCCACTGTCAAGCAACCACCTGCGGCCCTGCCCACTTTAGGTGCAACAGCAGCATCTG<br>CATCCCTAGCCTGTGGGCCTGCGATGGCGACGTGGACTGCGTGGATGGAAGCGATGAGTGGCCTCAGAACT<br>GCCAAGGAAGGGACACCGCTAGCAAGGGCGTGAGCAGCCCTTGCAGCAGCCTGGAGTTCCACTGCGGCAG<br>CAGCGAGTGCATCCATAGGAGCTGGGTCTGTGATGGTGAGGCGGATTGTAAGGACAAGAGCGACGAAGAAC<br>ATTGCGCCGTGGCCACCTGTAGGCCTGACGAATTTCAGTGCGCAGACGGCAGCTGCATCCACGGCTCTAGG<br>CAGTGCGATAGGGAGCACGACTGTAAGGATATGAGTGACGAGCTGGGCTGCGTGAACGTGACACAGTGCGA<br>CGGCCCTAACAAGTTCAAGTGCCACAGCGGCGAATGCATCAGCCTGGACAAGGTGTGCGACAGCGCTAGGG<br>ACTGCCAAGACTGGAGCGACGAGCCTATCAAGGAGTGCAAGACCAACGAGTGCCTGGACAACAACGGCGGC<br>TGCAGCCACATCTGCAAGGACCTGAAGATCGGCAGCGAGTGCCTGTGCCCTAGCGGCTTTAGGCTGGTGGA<br>CCTGCATAGGTGCGAGGACATCGACGAGTGCCAAGAGCCTGACACCTGCTCTCAGCTGTGCGTGAACCTGG<br>AGGGCAGCTACAAGTGCGAGTGCCAAGCCGGCTTCCACATGGACCCTCACACAAGGGTGTGCAAGGCCGTG<br>GGCAGCATCGGCTACCTGCTGTTCACCAATAGGCACGAGGTGAGGAAGATGACCCTGGATAGGAGCGAGTAC<br>ACAAGCTTACTGCCTAATCTGAAGAACGTGGTGGCCCTGGACACCGAGGTGACCAACAATAGGATCTACTGG<br>AGCGACCTGTCTCAGAAGAAGATCTACAGCGCCCTGATGGACCAAGCCCCTAACCTGAGCTACGACACCATC<br>ATCTCTGAAGACCTGCACGCCCCTGACGGCCTGGCCGTGGACTGGATCCATAGGAACATCTACTGGACTGAC<br>AGCGTGCCTGGCAGCGTGAGCGTGGCCGACACCAAGGGCGTGAAGAGGAGGACCCTGTTCCAAGAGGCCG<br>GCTCTAGGCCTAGGGCCATCGTGGTGGACCCTGTGCACGGCTTCATGTACTGGACCGATTGGGGCACCCCT<br>GCCAAGATCAAGAAGGGCGGCCTGAACGGCGTGGACATCCACAGCCTGGTGACCGAGAACATTCAGTGGCC<br>TAACGGCATCACCCTGGACCTGAGCAGCGGAAGGCTGTACTGGGTGGACAGCAAGCTGCACAGCATCAGCA<br>GCATCGACGTGAACGGCGGCAATAGGAAGACCATCCTGGAGGACGAAAATAGGCTGGCCCACCCTTTCAGC<br>CTGGCCATCTACGAGGACAAGGTGTACTGGACTGATGTGATTAACGAGGCCATCTTCAGCGCCAATAGGCTGA<br>CCGGCAGCGACGTGAACCTGGTGGCCGAGAACCTGCTGAGCCCTGAGGACATCGTGCTGTTCCACAAGGTG<br>ACACAGCCTAGGGGCGTGAACTGGTGCGAAACCACCGCCCTTCTGCCTAACAGCGGCTGTCAGTACCTGTG<br>CCTGCCTGCCCCTCAGATCGGCCCTCACAGCCCTAAGTTCACCTGCGCCTGCCCTGATGGCATGCTACTCGC<br>CGAGGACATGAGGAGCTGCCTGACGGAGGTGGACACCGTGCTGACCACTCAAGGCACCTCAGCAGTGAGG<br>CCTGTGGTGACCGCTAGCGCCACAAGGCCTCCTAAGCACTCTGAGGACCTCAGCGCCCCTAGCACCCCGAG<br>GCAGCCTGTGGACACCCCTGGCCTGAGCACCGTGGCCTCTGTGACCGTGAGCCACCAAGTGCAAGGCGAC<br>ATGGCGGGGAGGGGCAATGAGGAGCAGCCGCACGGCGTGAGGTTCCTGAGCATCTTCTTCCCTATCGCCCT<br>GGTGGCCCTGCTGGTGCTGGGCGCCGTGCTGCTGTGGAGGAACTGGAGGCTGAAGAACATCAACAGCATCA<br>ACTTCGACAACCCTGTGTATCAGAAGACCACCGAGGACGAGCTGCACATCTGTAGGAGCCAAGACGGCTACA<br>CCTACCCTTCTAGGCAGATGGTGAGCCTCGAAGACGACGTAGCCTGAgctcgctttcttgctgtcc |
| FWT7AGG | GCTCTAATACGACTCACTATAAGG |
| RVT7AGG | CCTTATAGTGAGTCGTATTAGAGC |
| FWhbb5UTR | AGCAACCTCAAACAGACACC |
| RVhbb5UTR | GGTGTCTGTTTGAGGTTGC |
| FWhbb3UTR | AGCTCGCTTTCTTGCTGTC |
| RVhbb3UTR | GCAATGAAAATAAATGTTTTTTATTAGGC |
| RVhbb3UTR_2 | GGACAGCAAGAAAGCGAGC |
| FWhbb3UTR_EGFP | GCTGTACAAGTAAGCTCGCTTTCTTGCTGTC |
| RVhbb5UTR_EGFP | CCTCGCCCTTGCTCACCATGGTGTCTGTTTGAGGTTG |
| RVhbb5UTR_GCCACC | GGTGGCTGTTTGAGGTTGC |
| RVhba5UTR | GGTGGCGGGTTCTCTCTG |
| RVhba5UTR_EGFP | CACCATGGTGGCGGGTTCTCTCTG |
| FWaes3UTR | CTGGTACTGCATGCACGC |
| FWaes3UTR_EGFP | GTACAAGTAACTGGTACTGCATGCACGC |
| FWmtRNRUTR | CAAGCACGCAGCAATG |
| RVmtRNRUTR | CATTGCTGCGTGCTTG |
| RVsynUTR | GGTGGCGGCACACTCG |
| RVsynUTR_2 | ACACTCGGAGTGCGG |
| RVsynUTR_GCC | GGCACACTCGGAGTG |
| RVsynUTR_GAC | GTCACACTCGGAGTGC |
| RVsynUTR_GACGCC | GGCGTCACACTCGGAGTGC |
| RVsynUTR_GCCGAC | GTCGGCACACTCGGAGTG |
| RVsynUTR_GACGAC | GTCGTCACACTCGGAGTG |
| RVS2UTR | GGTGTCGAATGGACGC |
| RVS2UTR_GCCACC | GGTGGCGAATGGACGC |
| RVS2UTR_GCCGACACC | GGCGAATGGACGCGTTATTTTG |
| RVden2_5UTR | CAGAGATCTGCTCTCTAATT |
| RVden2_5UTR_GAC | GTCCAGAGATCTGCTCTCTAATT |
| RVden2_GCC | GGCCAGAGATCTGCTCTCTAATT |

| | |
|---|---|
| RVden2_GCCGCC | GGCGGCCAGAGATCTGCTCTCTAATT |
| RVden2_GCCGAC | GTCGGCCAGAGATCTGCTCTCTAATT |
| RVden2_5UTR_GACGCC | GGCGTCCAGAGATCTGCTCTCTAATT |
| RVden2_5UTR_CCCGAC | GTCGGGCAGAGATCTGCTCTCTAATT |
| RVden2_5UTR_ACCGAG | CTCGGTCAGAGATCTGCTCTCTAATT |
| RVden2_5UTR_ACCGAC | GTCGGTCAGAGATCTGCTCTCTAATT |
| FWden2_5UTR_GCC | AATTAGAGAGCAGATCTCTGGCC |
| FWhbb5UTR_36nt | CACAACTGTGTTCACTAGCA |
| FWsynUTR_50nt | GTCTGTTCTGTTACACAAGGG |
| FWsyn2_5UTR_25nt | CAAAATAACGCGTCCATTCG |
| FWden2_5UTR_50nt | CTAAGCTCAACGTAGTTCTAACAG |
| FWden2_5UTR_25nt | TTTTTAATTAGAGAGCAGATCTCTG |
| RVT7AGG_GCCACC | GGTGGCCCTTATAGTGAGTCGTATTAGAGC |
| RVT7AGG_GCCGACACC | GGTGTCGGCCCTTATAGTGAGTCGTATTAGAGC |
| FW5pEGFP | ATGGTGAGCAAGGGCGAG |
| RV5pEGFP_2 | TCCTCGCCCTTGCTCAC |
| FW3pEGFP | CATGGACGAGCTGTACAAG |
| RV3pEGFP_AES3UTR | CGTGCATGCAGTACCAGTTACTTGTACAGCTCGTCCATG |
| FWegfp_HBA5UTR | CCCGCCACCATGGTGAGCAAGGGCGAG |
| RVegfp_mtRNRUTR | CTGCATTGCTGCGTGCTTGTTACTTGTACAGCTCGTCCATG |
| FWegfpCDS | CGCACCATCTTCTTCAAGG |
| RVegfpCDS | CCTTGAAGAAGATGGTGCG |
| FWFLuc | GCCTGAGAGCTTCGATAGG |
| RVFLuc | CCTATCGAAGCTCTCAGGC |
| FWFLuc_2 | CGAGCTGTGTGTTAGGGG |
| RVFLuc_2 | CCCCTAACACACAGCTCG |
| RV3pFLuc_AES3UTR | GCGTGCATGCAGTACCAGTCACACGGCGATCTTGC |
| RV3pFLuc_mtRNR3UTR | CTGCATTGCTGCGTGCTTGTCACACGGCGATCTTGC |
| FW5pFLuc | ATGGAGGACGCCAAGAAC |
| FW5pFLuc_HBA5UTR | CAGAGAGAACCCGCCACCATGGAGGACGCCAAGAAC |
| FW5pFLuc_SynUTR | GAGTGTGCCGCCGACACCATGGAGGACGCCAAGAAC |
| RV5pFLuc_Syn2UTR | CGCCGACACCATGGAGGACGCCAAGAAC |
| FW5pFLuc_den2GCC | AGCAGATCTCTGGCCACCATGGAGGACGCCA |
| FWLDLR | AGCCTGGAGTTCCACTGC |
| RVLDLR | CGCAGTGGAACTCCAGGC |
| FWLDLR_2 | CGACCTGTCTCAGAAGAAGA |
| RVLDLR_2 | TCTTCTTCTGAGACAGGTCG |
| FWLDLR_3 | TCAGATCGGCCCTCACAG |
| RVLDLR_3 | CTGTGAGGGCCGATCTGA |
| FW5pLDLR_HBA5UTR | CTCAGAGAGAACCCGCCACCATGAGCACCGCCG |
| RV3pLDLR_AES3UTR | TGCATGCAGTACCAGTCAGGCTACGTCGTCTTCG |
| FWpUTR_IVT_hbb_den2 | GTCCCAATGATGCTCTAATACG |
| RVhbb3UTR_pA120 | TTTTTTTTTTTTTTTTTTTTTTTTTTTTTTTTTTTTTTTTTTTTTTTTTTTTTTTTTTTTTTTTTTTTTTTTTTTTTTTTTT<br>TTTTTTTTTTTTTTTTTTTTTTTTTTTTTTTTTTTTTTGCAATGAAAATAAATGTTTTTTATTAGGC |
| RVden2_3UTR_pA120 | TTTTTTTTTTTTTTTTTTTTTTTTTTTTTTTTTTTTTTTTTTTTTTTTTTTTTTTTTTTTTTTTTTTTTTTTTTTTTTTTTT<br>TTTTTTTTTTTTTTTTTTTTTTTTTTTTTTTTTTTTTTTCTAGAACCTGTTGATTCAAC |
| FWpUTR _IVT_mtRNR | ACCTGACGTCCCAATGATGCTCTAATACG |
| RVmtRNR_pA120 | TTTTTTTTTTTTTTTTTTTTTTTTTTTTTTTTTTTTTTTTTTTTTTTTTTTTTTTTTTTTTTTTTTTTTTTTTTTTTTTTTT<br>TTTTTTTTTTTTTTTTTTTTTTTTTTTTTTTTTTTTTTGGTGTGGCTGGCACGAAATTGACCAAC |

**Table S4. *In vitro*-transcribed mRNAs and lipid nanoparticle formulations used in this study.**

| Formulation (lipid molar ratio) | Kozak / UTR variant (5'UTR(Kozak seq) / 3'UTR) | Hydrodynamic diameter (nm) | PDI | Zeta potential (mV) | Encapsulation efficiency (%) | mRNA # (Fig. S13) | Figures |
|---|---|---|---|---|---|---|---|
| **EGFP-mRNA formulations** | | | | | | | |
| **Kozak sequence variant screening** | | | | | | | |
| Dlin-MC3-DMA:DSPC:chol:DMG-PEG2K:DiD (50:10:38.3:1.5:0.2) | Syn1(no Kozak seq) / HBB | 103.20 ± 3.06 | 0.127 ± 0.034 | ND | 96.56 | 1 | 2, S1–2 |
| | Syn1(GCCACC) / HBB | 113.07 ± 0.55 | 0.146 ± 0.009 | ND | 96.09 | 2 | 2, S1–2 |
| | Syn1(GCCGCCACC) / HBB | 110.50 ± 1.30 | 0.118 ± 0.010 | ND | 97.82 | 3 | 2, S1–2 |
| | Syn1(GACACC) / HBB | 114.10 ± 2.95 | 0.108 ± 0.028 | ND | 96.82 | 4 | 2, S1–2 |
| | Syn1(GACGCCACC) / HBB | 92.95 ± 2.31 | 0.128 ± 0.022 | ND | 96.92 | 5 | 2, S1–2 |
| | Syn1(GCCGACACC) / HBB | 89.16 ± 0.90 | 0.100 ± 0.026 | ND | 99.99 | 6 | 2, S1–2 |
| | Syn1(GACGACACC) / HBB | 91.55 ± 2.13 | 0.120 ± 0.029 | ND | 97.84 | 7 | 2, S1–2 |
| | Den2(no Kozak seq) / HBB | 103.73 ± 1.53 | 0.095 ± 0.008 | ND | 94.45 | 8 | 2, S1–2 |
| | Den2(GCCACC) / HBB | 104.87 ± 2.11 | 0.115 ± 0.026 | ND | 96.60 | 9 | 2, S1–2 |
| | Den2(GCCGCCACC) / HBB | 114.70 ± 3.39 | 0.104 ± 0.021 | ND | 97.33 | 10 | 2, S1–2 |
| | Den2(GACACC) / HBB | 119.40 ± 3.86 | 0.091 ± 0.037 | ND | 97.96 | 11 | 2, S1–2 |
| | Den2(GACGCCACC) / HBB | 113.57 ± 2.73 | 0.121 ± 0.021 | ND | 97.16 | 12 | 2, S1–2 |
| | Den2(GCCGACACC) / HBB | 108.40 ± 0.95 | 0.171 ± 0.018 | ND | 96.92 | 13 | 2, S1–2 |
| | Den2(GACGACACC) / HBB | 103.37 ± 1.36 | 0.096 ± 0.009 | ND | 95.74 | 14 | 2, S1–2 |
| | No mRNA (empty control) | 115.20 ± 0.61 | 0.131 ± 0.007 | ND | N/A | N/A | S2 |
| **Initial combinatorial UTR optimization** | | | | | | | |
| Dlin-MC3-DMA:DSPC:chol:DMG-PEG2K:DiD (50:10:38.3:1.5:0.2) | HBB(GACACC) / HBB | 82.86 ± 1.31 | 0.088 ± 0.032 | -5.33 ± 1.06 | 97.71 | 15 | 3–4, S3–4 |
| | Syn1(GCCGCCACC) / HBB | 83.61 ± 1.21 | 0.149 ± 0.011 | -7.24 ± 0.64 | 97.77 | 16 | 3, S3–4 |
| | Syn2(GACACC) / HBB | 79.09 ± 2.26 | 0.109 ± 0.026 | -6.71 ± 0.49 | 99.81 | 17 | 3–4, S3–4 |
| | Den2(GCCGCCACC) / HBB | 86.22 ± 1.87 | 0.155 ± 0.015 | -6.63 ± 0.58 | 96.55 | 18 | 3, S3–4 |
| | HBB(GACACC) / mtRNR1 | 94.99 ± 1.95 | 0.107 ± 0.011 | -5.89 ± 0.23 | 96.30 | 19 | 3, S3–4 |
| | Syn1(GCCGCCACC) / mtRNR1 | 81.19 ± 0.33 | 0.104 ± 0.014 | -9.16 ± 0.99 | 98.57 | 20 | 3, S3–4 |
| | Syn2(GACACC) / mtRNR1 | 82.57 ± 0.37 | 0.096 ± 0.024 | -5.82 ± 1.01 | 97.88 | 21 | 3, S3–4 |
| | Den2(GCCGCCACC) / mtRNR1 | 86.40 ± 1.09 | 0.097 ± 0.038 | -6.83 ± 1.24 | 98.02 | 22 | 3, S3–4 |
| | HBB(GACACC) / AES-mtRNR1 | 85.88 ± 0.84 | 0.098 ± 0.008 | -6.00 ± 0.15 | 98.50 | 23 | 3, S3–4 |
| | Syn1(GCCGCCACC) / AES-mtRNR1 | 81.62 ± 0.26 | 0.119 ± 0.006 | -4.62 ± 0.80 | 98.37 | 24 | 3, S3–4 |
| | Syn2(GACACC) / AES-mtRNR1 | 88.03 ± 0.68 | 0.104 ± 0.014 | -8.57 ± 1.45 | 98.13 | 25 | 3, S3–4 |
| | Den2(GCCGCCACC) / AES-mtRNR1 | 98.66 ± 1.51 | 0.149 ± 0.021 | -6.34 ± 0.66 | 98.25 | 26 | 3, S3–4 |
| | HBA / AES-mtRNR1 (**BNT**) | 82.88 ± 2.61* | 0.130 ± 0.028* | -7.78 ± 1.79* | 96.81 ± 2.38* | 27 | 3–4, S3–4 |
| | Den2(GCCACC) / Den2 | 91.12 ± 0.40 | 0.129 ± 0.001 | -4.64 ± 0.32 | 99.23 | 28 | 3, S3–4 |
| | No 5'UTR(GCCACC) / HBB | 80.02 ± 1.39 | 0.132 ± 0.017 | -4.05 ± 0.60 | 97.03 | 29 | 3, S3–4 |
| | No 5'UTR(GCCGACACC) / HBB | 105.43 ± 0.74 | 0.186 ± 0.007 | -5.79 ± 0.91 | 96.60 | 30 | 3, S3–4 |
| | Commercial EGFP-mRNA (**TriLink**) | 90.89 ± 1.03* | 0.156 ± 0.015* | -5.95 ± 0.24* | 97.81 ± 0.39* | N/A | 3–4, S3–4 |
| **Second combinatorial screening** | | | | | | | |
| Dlin-MC3-DMA:DSPC:chol:DMG-PEG2K:DiD (50:10:38.3:1.5:0.2) | HBB(G**C**CACC) / HBB | 94.06 ± 0.51 | 0.176 ± 0.006 | -9.13 ± 0.33 | 95.28 | 31 | 4, S5–6 |
| | Syn2(G**C**CACC) / HBB | 92.27 ± 1.06 | 0.145 ± 0.024 | -7.26 ± 0.54 | 98.18 | 32 | 4, S5–6 |
| | Syn2(GCCG**A**CACC) / HBB | 108.33 ± 1.01 | 0.180 ± 0.003 | -6.95 ± 0.83 | 98.90 | 33 | 4, S5–6 |
| | **25 nt**-Syn2(GCCG**A**CACC) / HBB | N/A (only tested *in vitro*) | | | | 34 | 4, S5 |
| | Den2(G**C**CACC) / HBB | 106.47 ± 6.59 | 0.170 ± 0.009 | -5.54 ± 0.82 | 96.80 | 35 | 4, S5–6 |
| | Den2(G**A**CACC) / HBB | 96.74 ± 1.23 | 0.179 ± 0.010 | -5.03 ± 1.99 | 93.89 | 36 | 4, S5–6 |
| | Den2(GCCG**A**CACC) / HBB | 85.29 ± 1.90 | 0.136 ± 0.019 | -6.09 ± 1.98 | 95.15 | 37 | 4, S5–6 |
| | **50 nt**-Den2(GCCG**A**CACC) / HBB | 79.15 ± 0.65 | 0.141 ± 0.018 | -7.20 ± 0.41 | 94.04 | 38 | 4, S5–6 |
| | **25 nt**-Den2(GCCG**A**CACC) / HBB | 96.03 ± 2.42 | 0.159 ± 0.011 | -6.89 ± 0.45 | 96.59 | 39 | 4, S5–6 |
| | Den2(GCCGACACC) / mtRNR1 | N/A (only tested *in vitro*) | | | | 40 | 4, S5 |
| | Den2(GCCGACACC) / AES-mtRNR1 | N/A (only tested *in vitro*) | | | | 41 | 4, S5 |
| | Den2(**A**CCG**A**CACC) / HBB | 112.40 ± 1.15 | 0.114 ± 0.004 | -2.14 ± 0.35 | 98.74 | 42 | S7–8 |
| | Den2(**C**CCG**A**CACC) / HBB | 116.37 ± 2.70 | 0.113 ± 0.011 | -1.19 ± 0.59 | 96.18 | 43 | S7–8 |
| | Den2(**A**CCG**AG**ACC) / HBB | 122.50 ± 2.97 | 0.104 ± 0.013 | -7.13 ± 2.12 | 94.76 | 44 | S7–8 |
| **Mice experiments** | | | | | | | |
| **FLuc-mRNA formulations** | | | | | | | |
| SM-102:DSPC:chol:DMG-PEG2K: DSPE-PEG2K-triGalNAc (50:10:38.45:1.5:0.05) | No 5'UTR(GCCG**A**CACC) / HBB | 91.86 ± 3.69* | 0.127 ± 0.030* | -2.63 ± 0.92 | 94.89* | 45 | 5, S9 |
| | Syn1(GCCG**A**CACC) / HBB | 93.08 ± 3.20* | 0.135 ± 0.020* | -2.22 ± 0.75 | 94.75* | 46 | 5, S9 |
| | Syn2(GCCG**A**CACC) / HBB | 81.44 ± 7.35* | 0.130 ± 0.033* | -3.30 ± 0.43 | 96.00* | 47 | 5, S9 |
| | Den2(G**C**CACC) / HBB | 92.00 ± 0.48 | 0.154 ± 0.021 | -1.87 ± 1.09 | 95.27 | 48 | 5, S9 |
| | Den2(GCCG**A**CACC) / HBB | 80.19 ± 2.81* | 0.168 ± 0.018* | -1.18 ± 0.41 | 96.85* | 49 | 5, S9 |
| | **50 nt**-Den2(GCCG**A**CACC) / HBB | 92.35 ± 0.81 | 0.150 ± 0.012 | -5.58 ± 0.47 | 96.20 | 50 | 5, S9 |
| | **25 nt**-Den2(GCCG**A**CACC) / HBB | 86.81 ± 6.15* | 0.134 ± 0.018* | -3.05 ± 0.69 | 94.05* | 51 | 5, S9 |
| | Den2(GCCG**A**CACC) / AES-mtRNR1 | 97.93 ± 1.36 | 0.139 ± 0.018 | -1.54 ± 0.47 | 95.30* | 52 | 5, S9 |
| | **25 nt**-Den2(GCCG**A**CACC) / AES-mtRNR1 | 85.73 ± 1.16 | 0.097 ± 0.011 | -1.67 ± 0.13 | 95.27 | 53 | 5, S9 |
| | HBA / AES-mtRNR1 (**BNT**) | 97.43 ± 1.32 | 0.100 ± 0.013 | -3.07 ± 0.35 | 94.36* | 54 | 5, S9 |
| **Ldlr-mRNA formulations** | | | | | | | |
| SM-102:DSPC:chol:DMG-PEG2K: DSPE-PEG2K-triGalNAc (50:10:38.45:1.5:0.05) | **25 nt**-Den2(GCCG**A**CACC) / HBB | 99.08 ± 8.86* | 0.172 ± 0.010* | -5.14 ± 1.79* | 95.17 ± 0.40* | 55 | 5 |
| | Den2(GCCG**A**CACC) / AES-mtRNR1 | 96.02 ± 4.72* | 0.178 ± 0.048* | -4.69 ± 0.64* | 94.03 ± 2.49* | 56 | 5 |
| | HBA / AES-mtRNR1 (**BNT**) | 95.62 ± 0.71* | 0.152 ± 0.026* | -3.96 ± 1.64* | 95.07 ± 0.29* | 57 | 5 |

*Combined data of 2 independent LNP batches with mRNA from the same batch

**Table S5. Kozak sequence variants ranked by relative protein expression within each model system.** Expression levels were normalized within each 5′ UTR context to the highest value. Only the HBB 3′ UTR at 500 ng mL$^{-1}$ (*in vitro*) was used for this ranking. Across experiments with varying mRNA doses in zebrafish embryos, the GCCGACACC group was always included and performed best. Thus, that group served as the normalization reference.

| –9 to –1 position relative to start codon | *in vitro* | | | | *in vivo* | | | |
|---|---|---|---|---|---|---|---|---|
| **Kozak sequence** | **DC2.4** | **HEK293T** | **HeLa** | **HUVECs** | **Zebrafish** | **Mice_6hpi** | **Mice_24hpi** | **5′ UTR** |
| (ACA)GCCACC | 71,74 | 84,84 | 88,84 | 94,35 | 59,67 | ND | ND | HBB |
| (ACA)GACACC | 100,00 | 100,00 | 100,00 | 100,00 | 100,00 | ND | ND | HBB |
| (UGU)GCCACC | 41,46 | 59,79 | 37,52 | 57,39 | 55,52 | ND | ND | Syn1 |
| GCCGCCACC | 54,25 | 62,63 | 54,81 | 84,88 | 64,47 | ND | ND | Syn1 |
| (UGU)GACACC | 49,42 | 80,67 | 88,35 | 100,00 | 100,00 | ND | ND | Syn1 |
| GACGCCACC | 93,11 | 64,20 | 92,49 | 76,05 | 79,10 | ND | ND | Syn1 |
| GCCGACACC | 100,00 | 100,00 | 100,00 | 87,38 | 94,34 | 72,62 | ND | Syn1 |
| GACGACACC | 24,80 | 20,58 | 47,18 | 88,69 | 57,93 | ND | ND | Syn1 |
| (UUC)GCCACC | 41,04 | 60,02 | 64,47 | 100,00 | 37,82 | ND | ND | Syn2 |
| (UUC)GACACC | 64,91 | 100,00 | 95,96 | 95,17 | 100,00 | ND | ND | Syn2 |
| GCCGACACC | 100,00 | 73,79 | 100,00 | 95,35 | 73,68 | 55,57 | ND | Syn2 |
| (CUG)GCCACC | 34,79 | 71,86 | 51,78 | 19,17 | 58,38 | 27,59 | 49,86 | Den2 |
| GCCGCCACC | 58,69 | 91,31 | 58,64 | 92,62 | 62,90 | ND | ND | Den2 |
| (CUG)GACACC | 66,70 | 91,98 | 59,06 | 84,46 | 76,51 | ND | ND | Den2 |
| GACGCCACC | 62,89 | 34,39 | 44,05 | 70,28 | 50,96 | ND | ND | Den2 |
| GCCGACACC | 96,00 | 100,00 | 100,00 | 100,00 | 100,00 | 100,00 | 100,00 | Den2 |
| GACGACACC | 65,27 | 43,31 | 60,48 | 63,78 | 45,76 | ND | ND | Den2 |
| ACCGACACC | 100,00 | ND | ND | 89,46 | 82,54 | ND | ND | Den2 |
| CCCGACACC | 93,12 | ND | ND | 91,66 | 62,91 | ND | ND | Den2 |
| ACCGAGACC | 83,69 | ND | ND | 69,41 | 97,82 | ND | ND | Den2 |

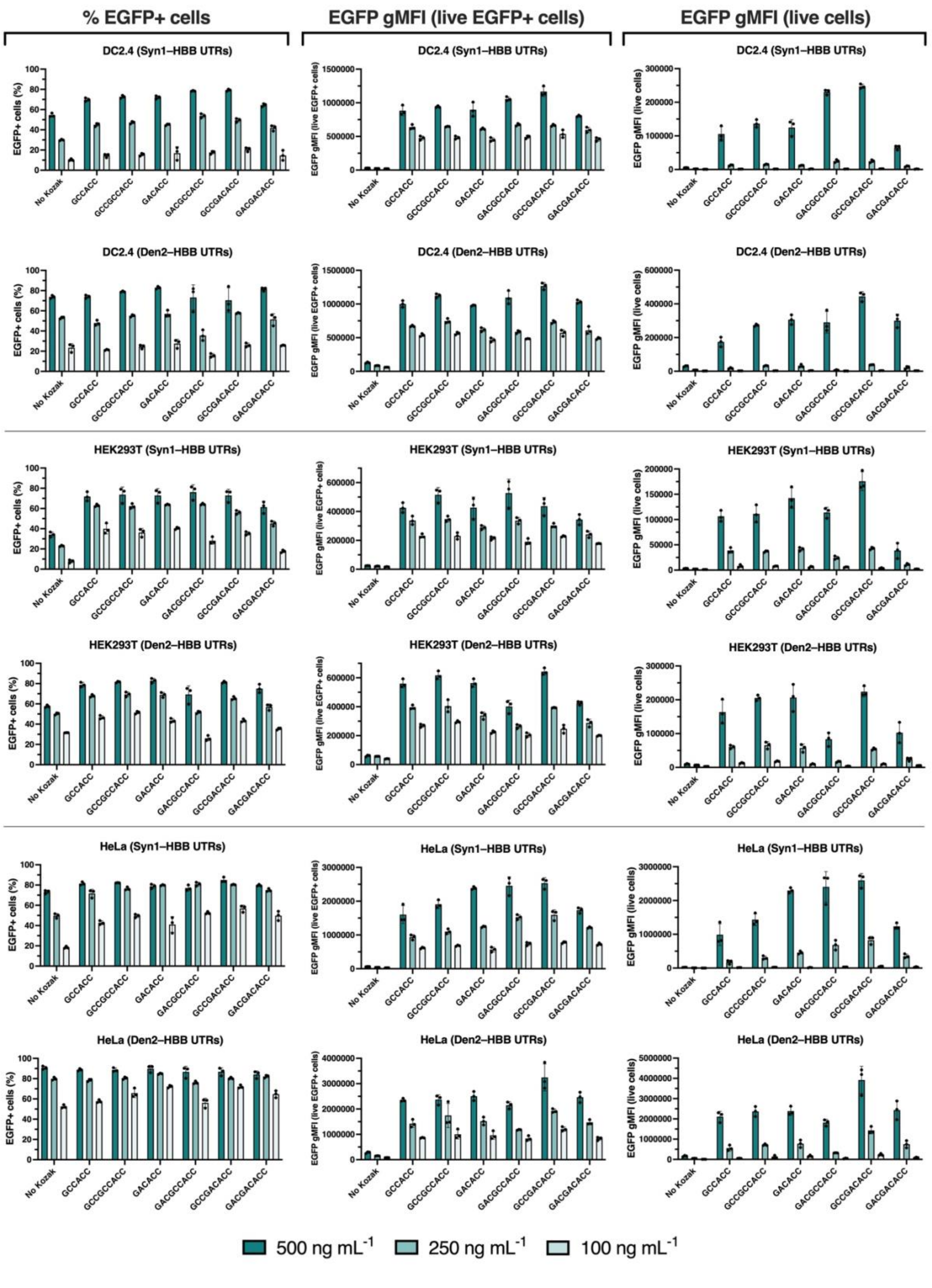

% EGFP+ cells
EGFP gMFI (live EGFP+ cells)
EGFP gMFI (live cells)
DC2.4 (Syn1–HBB UTRs)
DC2.4 (Den2–HBB UTRs)
HEK293T (Syn1–HBB UTRs)
HEK293T (Den2–HBB UTRs)
HeLa (Syn1–HBB UTRs)
HeLa (Den2–HBB UTRs)
EGFP+ cells (%)
EGFP gMFI (live EGFP+ cells)
EGFP gMFI (live cells)
No Kozak
GCCACC
GCCGCCACC
GACACC
GACGCCACC
GCCGACACC
GACGACACC
500 ng mL$^{-1}$
250 ng mL$^{-1}$
100 ng mL$^{-1}$

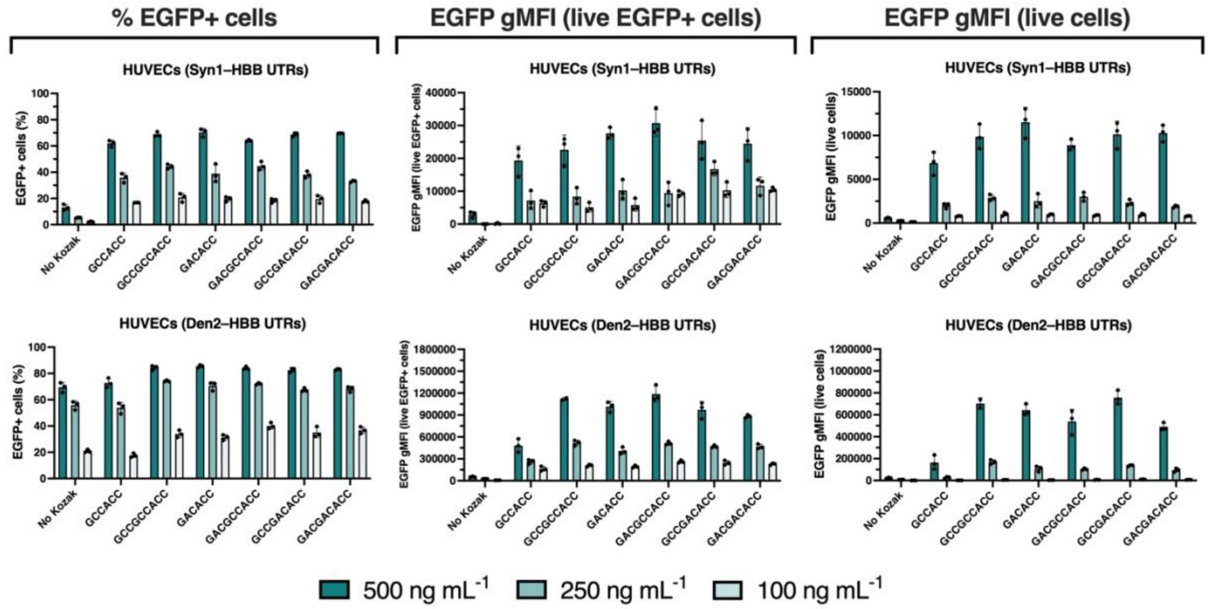


**Figure S1. Influence of Kozak sequence variation on EGFP expression *in vitro*.** DC2.4, HEK293T, HeLa, and HUVECs received 100–500 ng mL$^{-1}$ IVT mRNAs in lipoplexes with either the Syn1 (NeoUTR3) or Den2 5′ UTR and the HBB 3′ UTR carrying different Kozak sequence variants. 24 h later, cells were harvested, and EGFP expression was determined using flow cytometry, enabling the determination of % EGFP positive cells, EGFP geometric mean fluorescence intensity (gMFI) within the positive population, and EGFP gMFI of all cells.

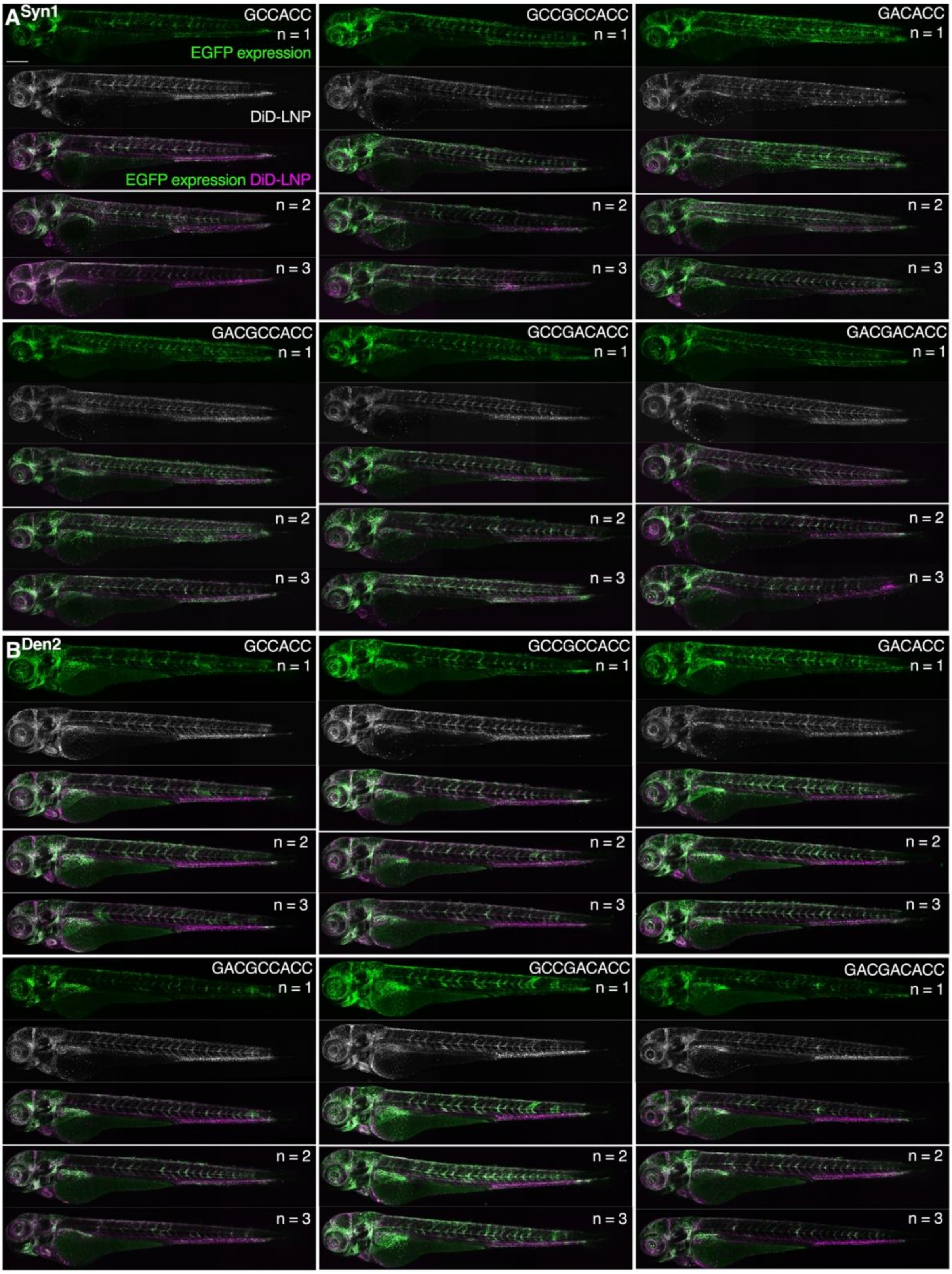
A Syn1
GCCACC
n = 1
EGFP expression
DiD-LNP
EGFP expression DiD-LNP
n = 2
n = 3
GCCGCCACC
GACACC
GACGCCACC
GCCGACACC
GACGACACC
B Den2

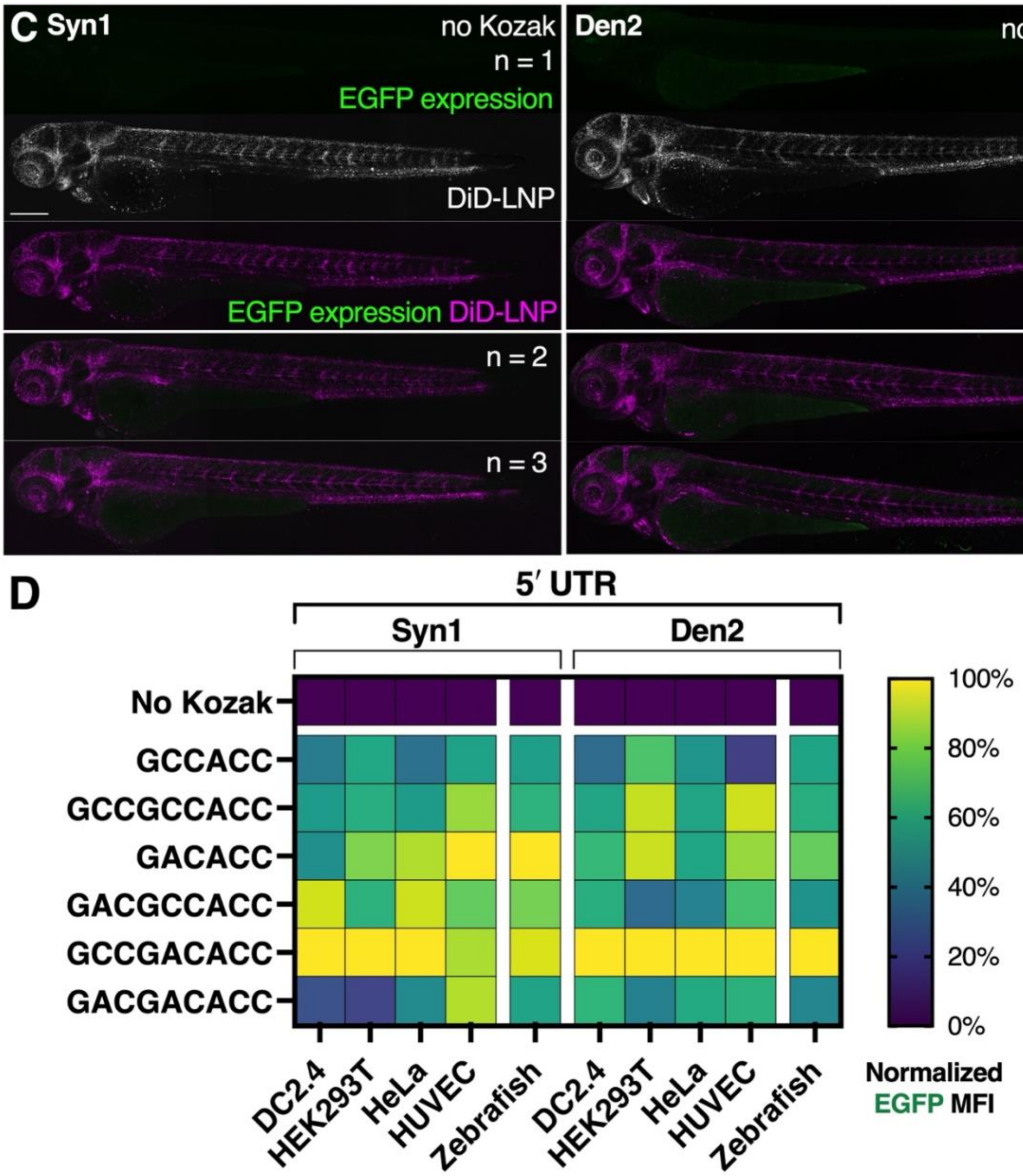


**Figure S2. Influence of Kozak sequence variation on EGFP expression in zebrafish embryos. A–C)** Confocal maximum z-projections showing EGFP expression (green) in whole zebrafish embryos 24 h after receiving (i.v.) 0.2 mg/kg mRNA encapsulated in DiD-labeled Onpattro-like LNPs (magenta) with either the Syn1 (NeoUTR3) (**A**) or Den2 5′ UTR (**B**) carrying different Kozak sequence variants, or without a Kozak sequence (**C**) and the HBB 3′ UTR. Scale bars represent 250 µm. **D)** Heatmap showing relative EGFP expression levels, normalized to the highest group per cell line and in zebrafish embryos, 24 h after lipofection with 500 ng mL$^{-1}$ or i.v. injection of 0.2 mg/kg EGFP-mRNA carrying Kozak sequence variants of either the Syn1 or Den2 5′ UTR paired with the HBB 3′ UTR.

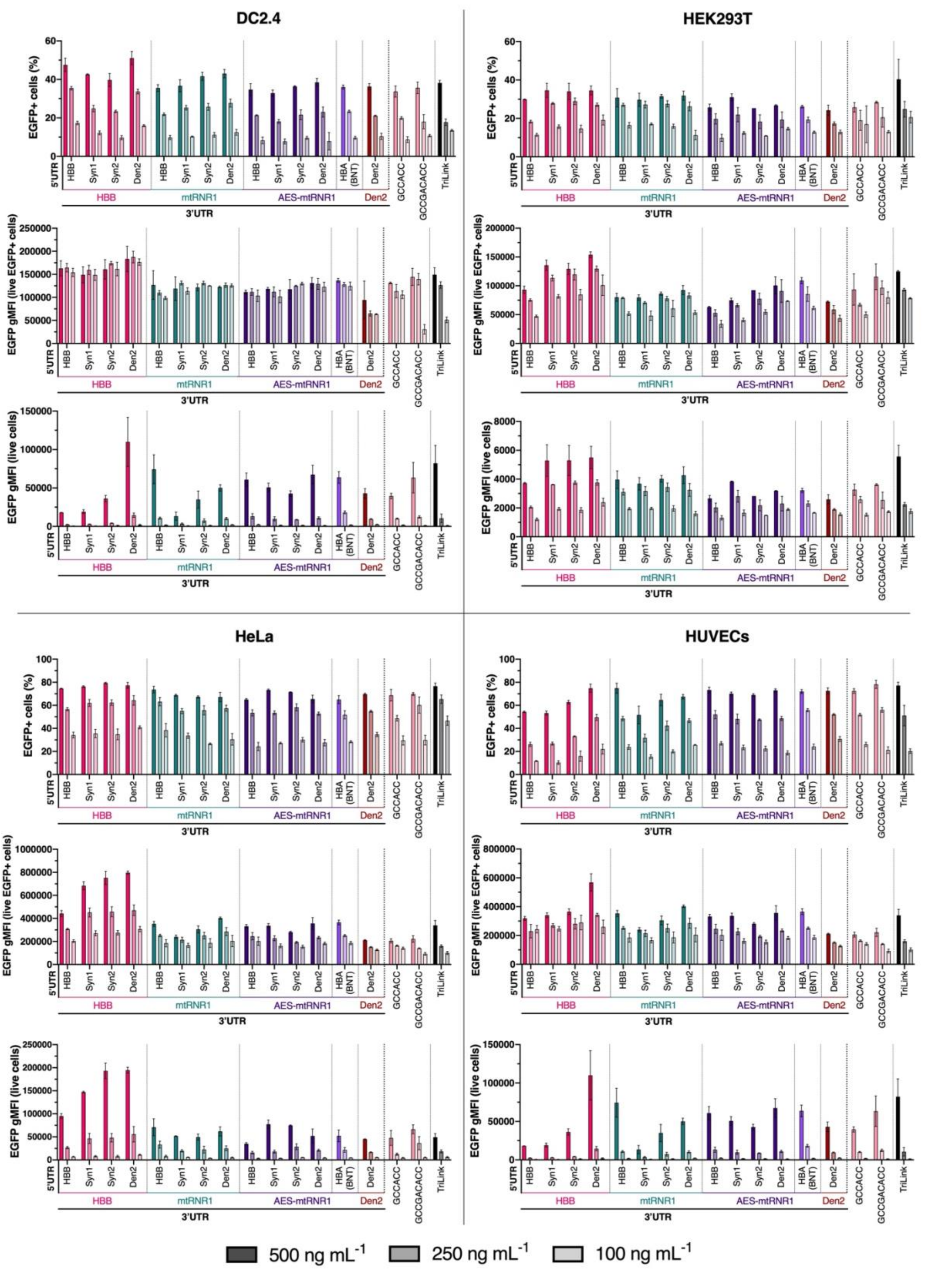


**Figure S3. Combinatorial UTR optimization *in vitro*.** DC2.4, HEK293T, HeLa, and HUVECs received 100–500 ng mL$^{-1}$ IVT mRNAs in lipoplexes carrying various UTR combinations, references with Kozak sequences lacking a 5′ UTR, or commercial EGFP-mRNA. 24 h later, cells were harvested, and EGFP expression was determined using flow cytometry, enabling the determination of % EGFP positive cells, EGFP geometric mean fluorescence intensity (gMFI) within the positive population, and EGFP gMFI of all cells.

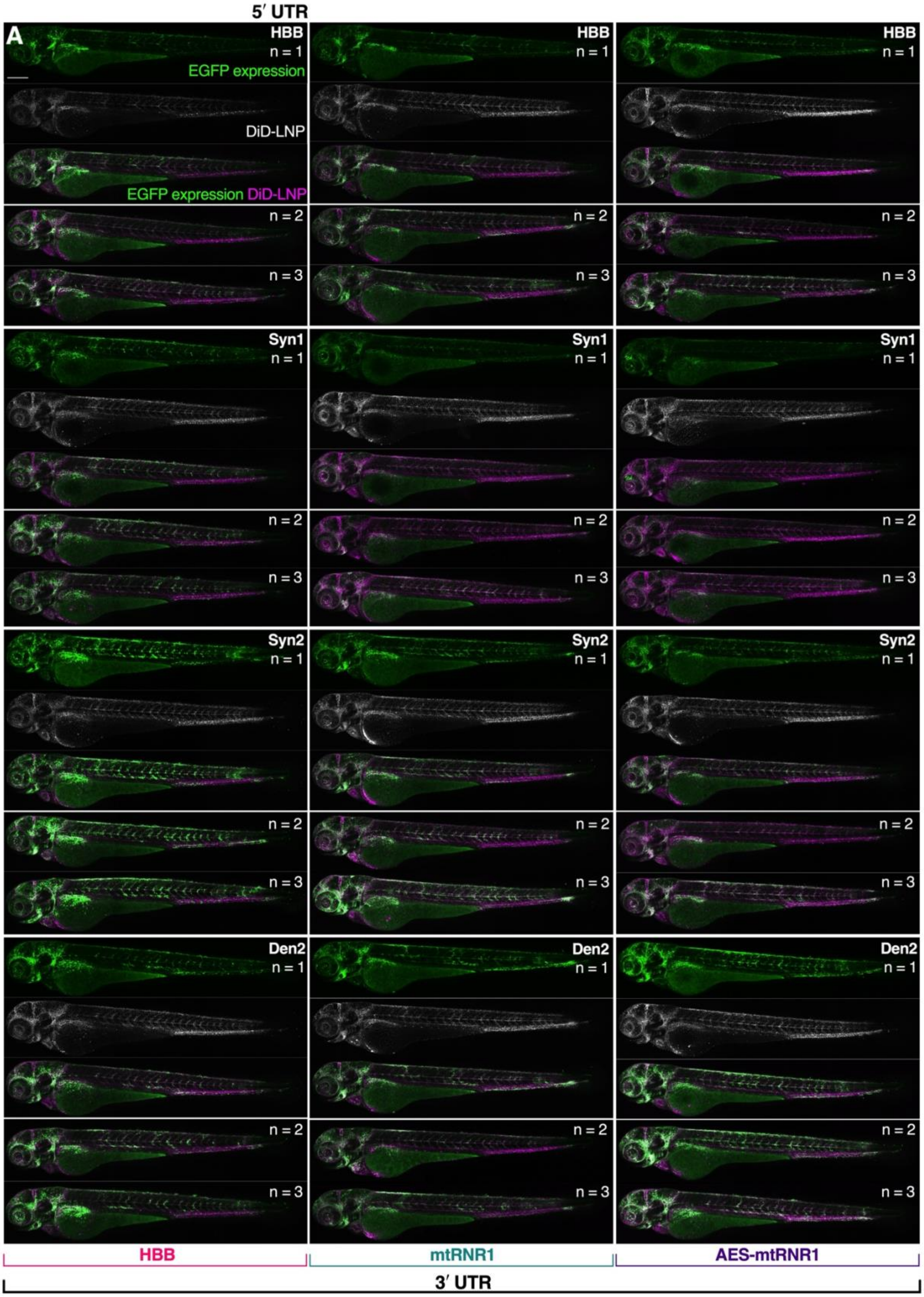

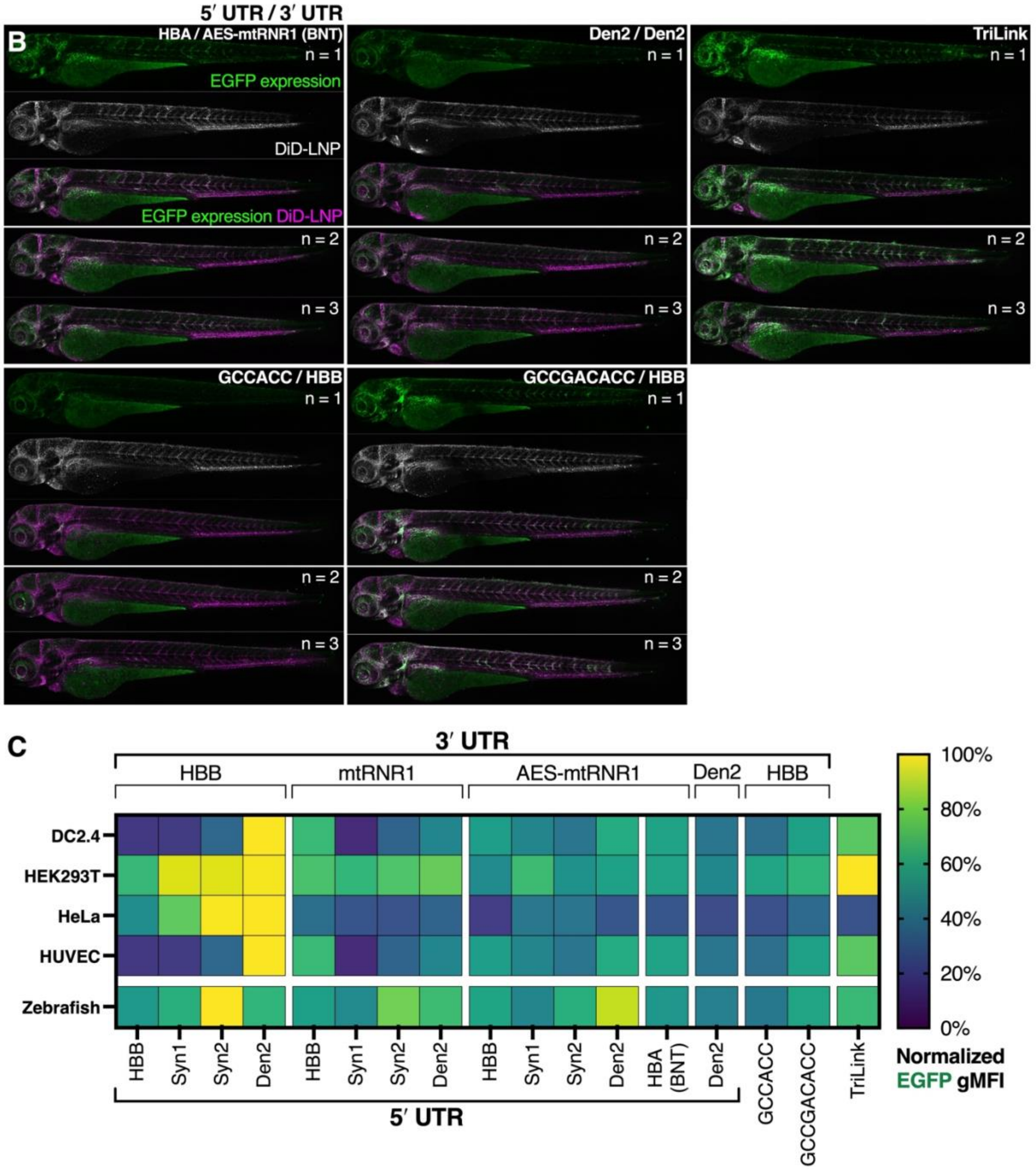


**Figure S4. Combinatorial UTR optimization in zebrafish embryos. A–B)** Confocal maximum z-projections showing EGFP expression (green) in whole zebrafish embryos 24 h after receiving (i.v.) 0.2 mg/kg IVT mRNA encapsulated in DiD-labeled Onpattro-like LNPs (magenta) carrying various UTR combinations (**A**), or references: with Kozak sequences lacking a 5′ UTR, Den2 UTRs, BNT UTRs, or commercial EGFP-mRNA (**B**). Scale bars represent 250 µm. **C)** Heatmap showing relative EGFP expression levels, normalized to the highest group per cell line and in zebrafish embryos, 24 h after lipofection with 500 ng mL$^{-1}$ or i.v. injection of 0.2 mg/kg EGFP-mRNA carrying various UTR combinations.

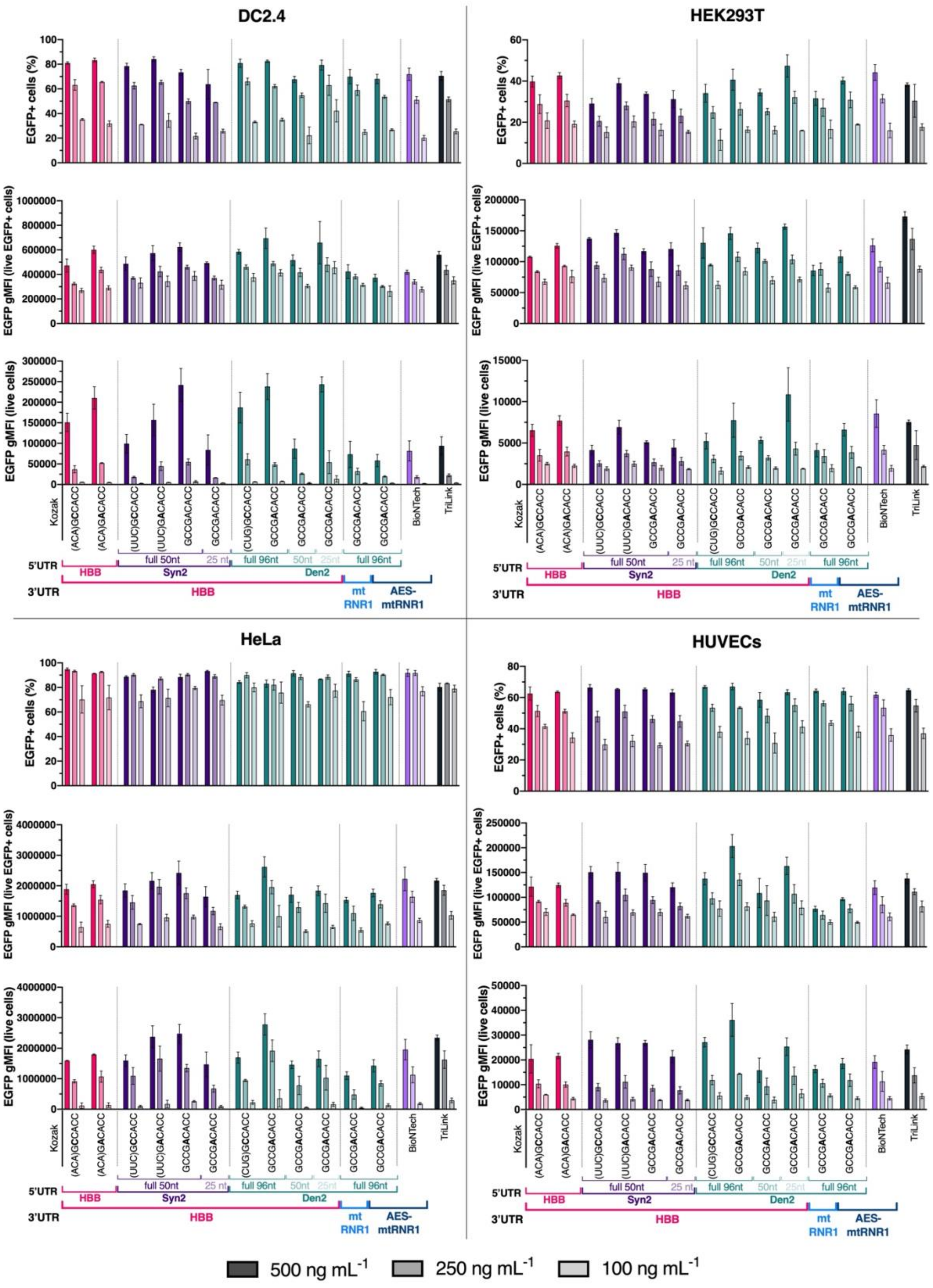


**Figure S5. Combined effect of a G-to-A substitution at the –5 position and 5′ UTR truncation on EGFP expression *in vitro*.** DC2.4, HEK293T, HeLa, and HUVECs received 100–500 ng mL$^{-1}$ IVT mRNAs in lipoplexes carrying different Kozak sequences and (truncated) UTR variants. 24 h later, cells were harvested, and EGFP expression was determined using flow cytometry, enabling the determination of % EGFP positive cells, EGFP geometric mean fluorescence intensity (gMFI) within the positive population, and EGFP gMFI of all cells.

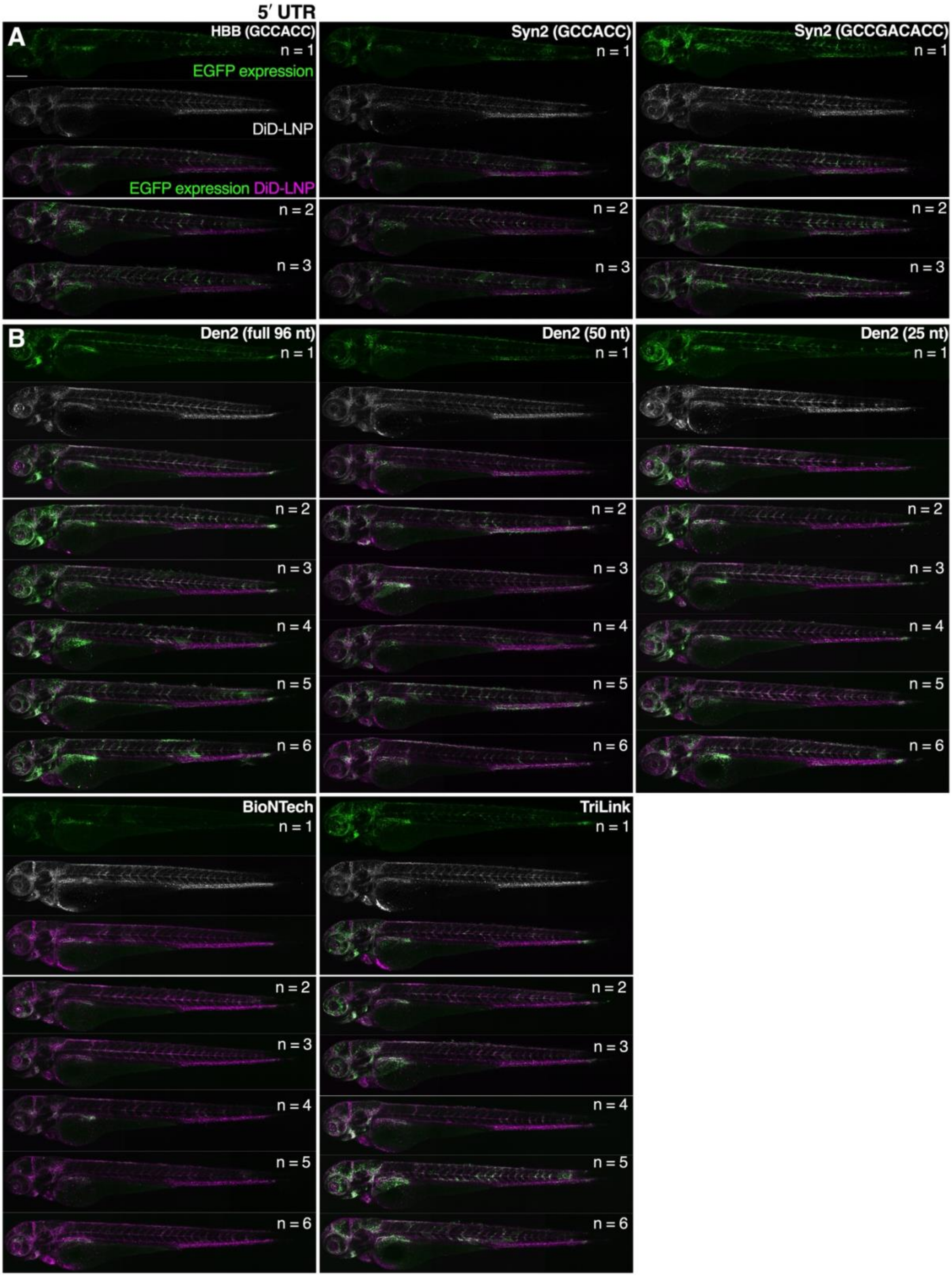


**Figure S6. Combined effect of a G-to-A substitution at the –5 position and 5′ UTR truncation on EGFP expression in zebrafish embryos. A)** Confocal maximum z-projections showing EGFP expression (green) in whole zebrafish embryos 24 h after receiving (i.v.) 0.2 mg/kg IVT mRNA encapsulated in DiD-labeled

Onpattro-like LNPs (magenta) carrying HBB or Syn2 with A-to-G substitutions at the −5 position. These 5′ UTRs with their original Kozak sequences were evaluated in a previous experiment (Figures 3, S5) using the same microscope settings and lipid batches. **B)** Confocal maximum z-projections showing EGFP expression in whole zebrafish embryos 24 h after receiving (i.v.) 0.12 mg/kg IVT mRNA encapsulated in DiD-labeled Onpattro-like LNPs carrying full-length or truncated Den2 5′ UTR variants or reference EGFP-mRNAs. Scale bars represent 250 µm.

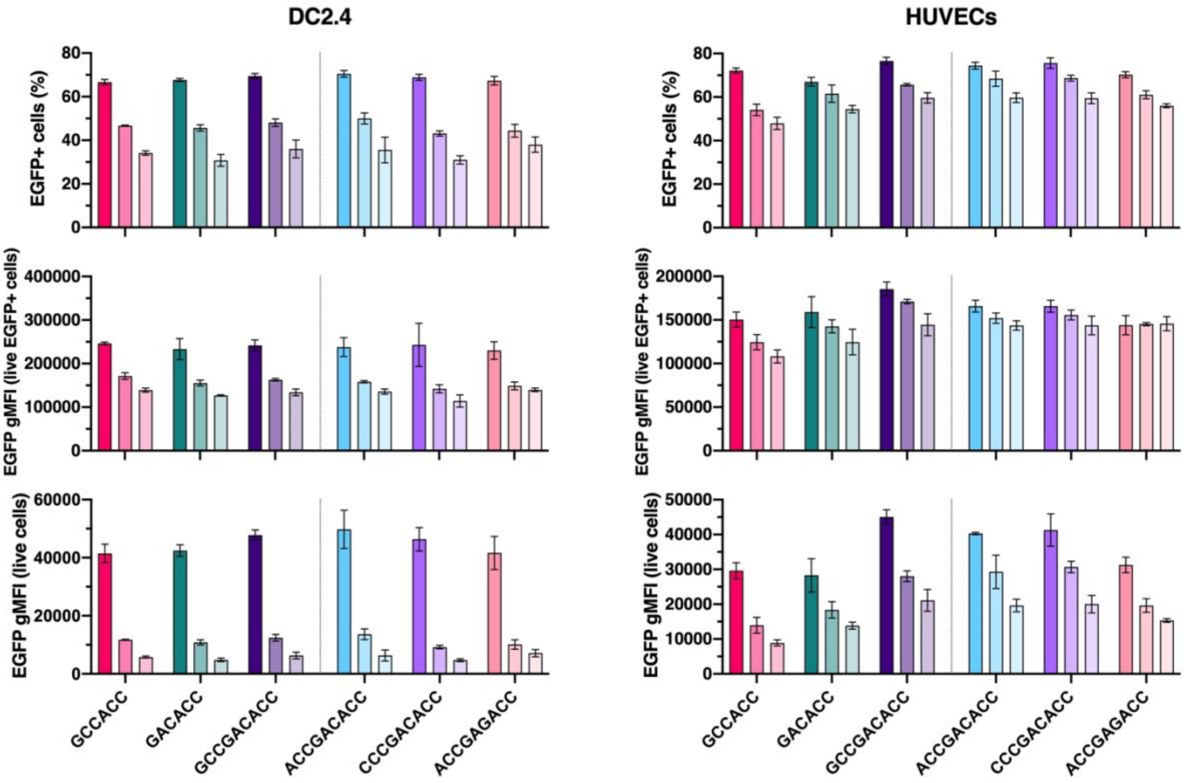


**Figure S7. Evaluation of additional Kozak sequence variants with Den2 5′ UTR *in vitro*.** DC2.4, and HUVECs received 100–500 ng mL$^{-1}$ IVT mRNAs in lipoplexes carrying different Kozak sequences differing at the −9 position or previously optimized for retinal genes (ACCGAGACC) with the Den2 5′ UTR and HBB 3′ UTR. 24 h later, cells were harvested, and EGFP expression was determined using flow cytometry, enabling the determination of % EGFP positive cells, EGFP geometric mean fluorescence intensity (gMFI) within the positive population, and EGFP gMFI of all cells.

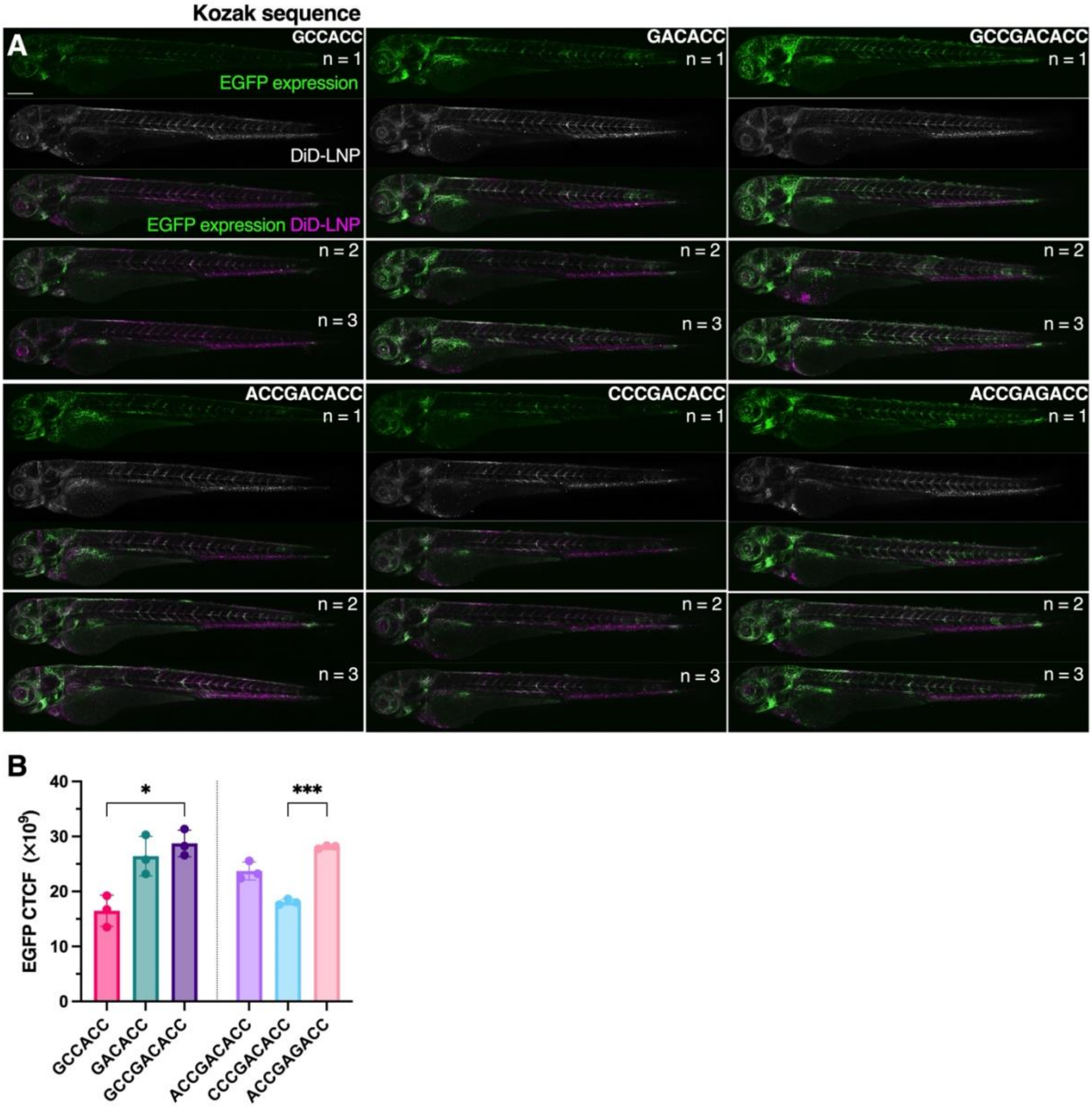


**Figure S8. Evaluation of additional Kozak sequence variants with Den2 5′ UTR in zebrafish embryos.** **A)** Confocal maximum z-projections showing EGFP expression (green) in whole zebrafish embryos 24 h after receiving (i.v.) 0.12 mg/kg IVT mRNA encapsulated in DiD-labeled Onpattro-like LNPs (magenta) carrying different Kozak sequences differing at the −9 position or previously optimized for retinal genes (ACCGAGACC) with the Den2 5′ UTR and HBB 3′ UTR. Scale bars represent 250 μm. **B)** Quantification of EGFP expression levels in wild-type zebrafish embryos that received Kozak sequence variants by corrected total cell fluorescence (CTCF). Data are presented as group means (n=3) ± SD and are statistically compared by a Brown-Forsythe & Welch ANOVA with Dunnett T3 correction (* p>0.05, *** p>0.001).

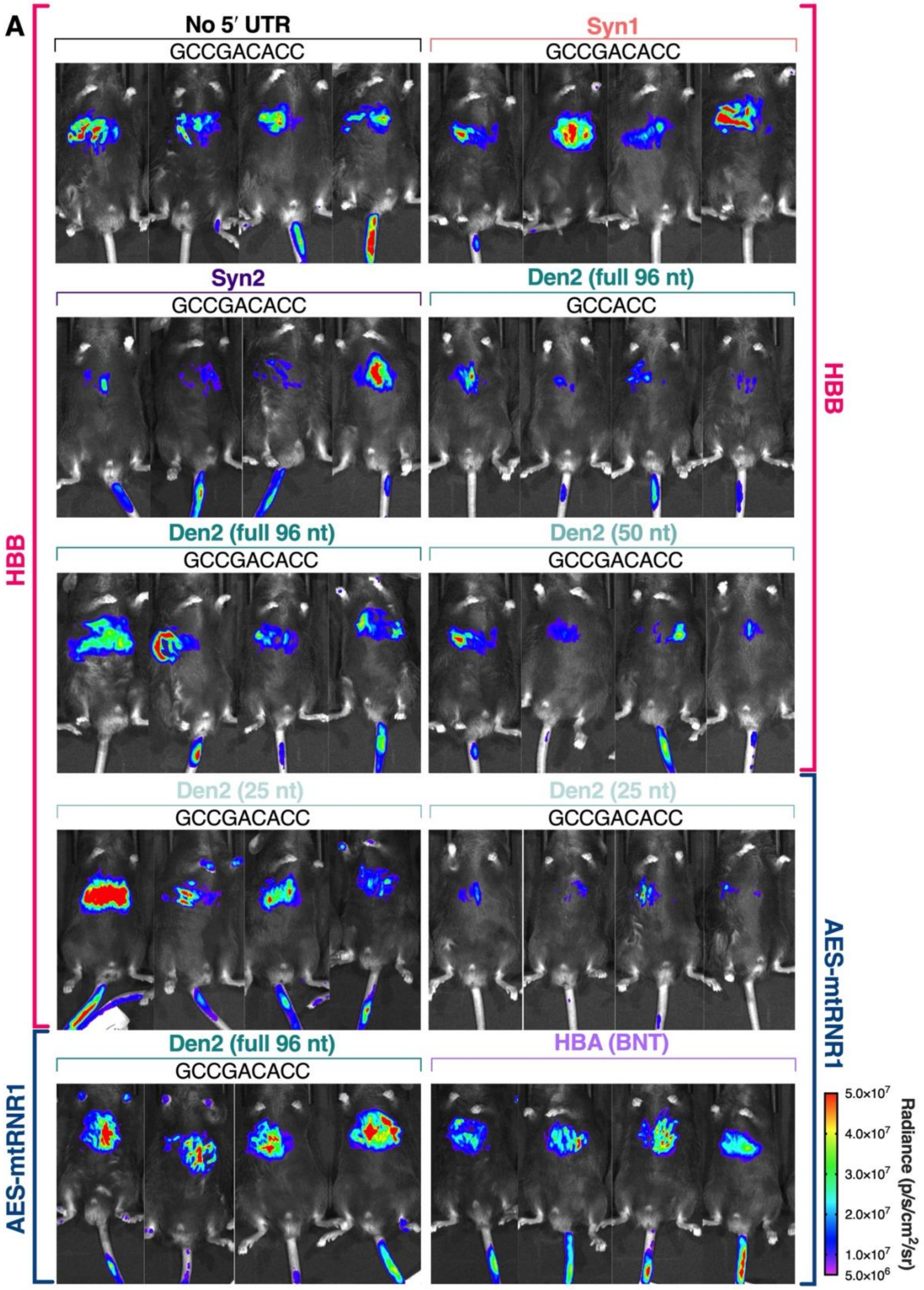
A
No 5′ UTR
GCCGACACC
Syn1
GCCGACACC
Syn2
GCCGACACC
Den2 (full 96 nt)
GCCACC
Den2 (full 96 nt)
GCCGACACC
Den2 (50 nt)
GCCGACACC
Den2 (25 nt)
GCCGACACC
Den2 (25 nt)
GCCGACACC
Den2 (full 96 nt)
GCCGACACC
HBA (BNT)
HBB
HBB
AES-mtRNR1
AES-mtRNR1
Radiance (p/s/cm²/sr)
5.0x10⁷
4.0x10⁷
3.0x10⁷
2.0x10⁷
1.0x10⁷
5.0x10⁶

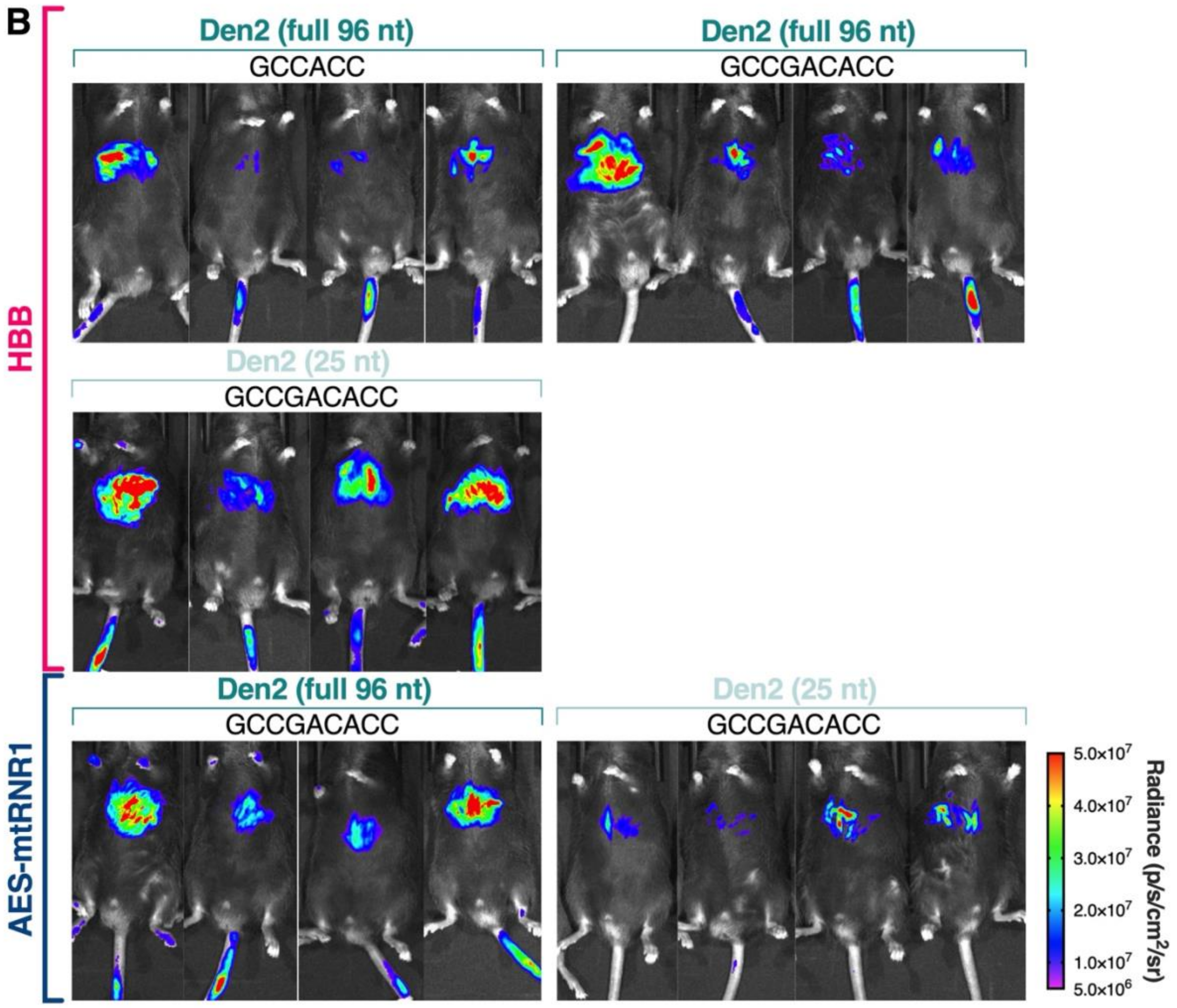


**Figure S9. Influence of different UTR or Kozak sequence combinations on luciferase expression in mice. A)** Live bioluminescence IVIS images of all *Ldlr* $^{-/-}$ mice 6 h after receiving 0.15 mg/kg FLuc-mRNA carrying various UTR or Kozak sequence combinations encapsulated in triGalNAc-functionalized LNPs. **B)** Live bioluminescence IVIS images of all *Ldlr* $^{-/-}$ mice 24 h after receiving 0.15 mg/kg FLuc-mRNA encapsulated in triGalNAc-functionalized LNPs.

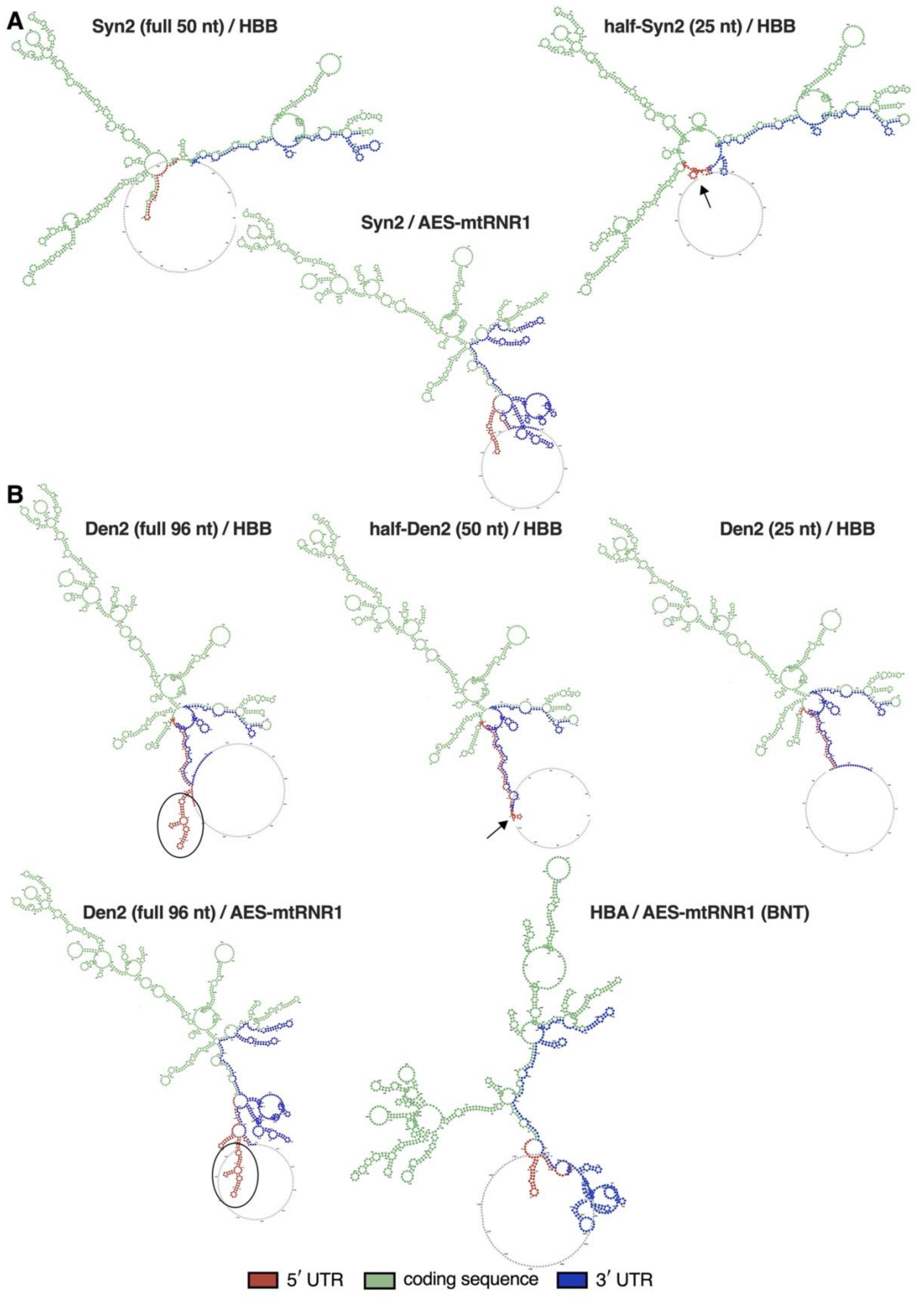

A
Syn2 (full 50 nt) / HBB
half-Syn2 (25 nt) / HBB
Syn2 / AES-mtRNR1
B
Den2 (full 96 nt) / HBB
half-Den2 (50 nt) / HBB
Den2 (25 nt) / HBB
Den2 (full 96 nt) / AES-mtRNR1
HBA / AES-mtRNR1 (BNT)
5′ UTR
coding sequence
3′ UTR

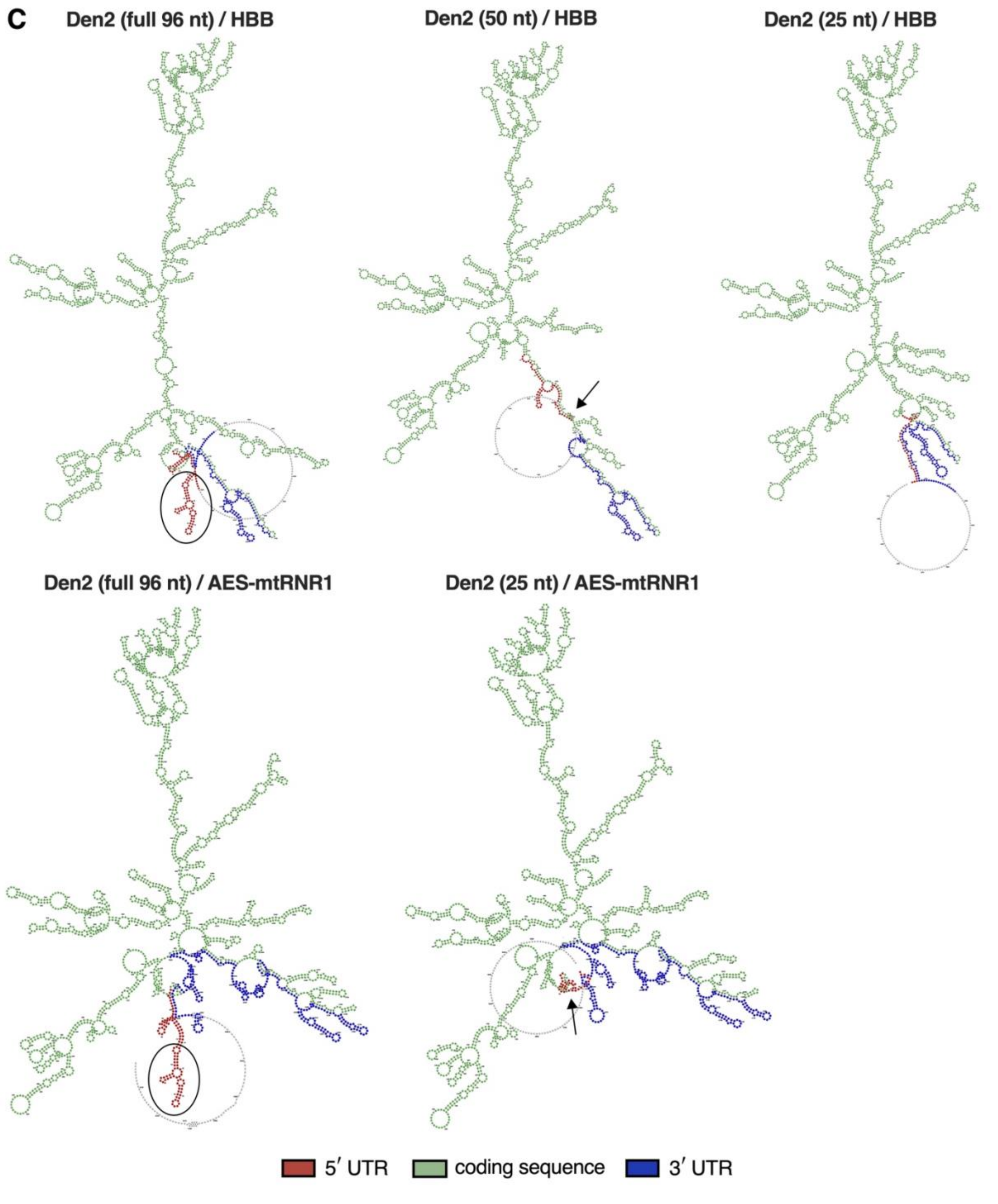


**Figure S10. mRNA secondary structure predictions comparing 5′ UTR truncation and coding sequence effects. A, B)** RNAfold-based structure prediction of EGFP-mRNAs carrying different (truncated) UTR combinations: Syn2 (**A**), and Den2 (**B**) 5′ UTR variants with the BNT UTRs as reference. **C)** RNAfold-based structure prediction of FLuc-mRNAs carrying (truncated) Den2 5′ UTR combinations. Encircled stem–loop regions in full-length Den2 variants (**B, C**) indicate stable structures that are not altered by paired 3′ UTR or CDS. Black arrows indicate highly structured, possibly less accessible regions in truncated 5′ UTRs.

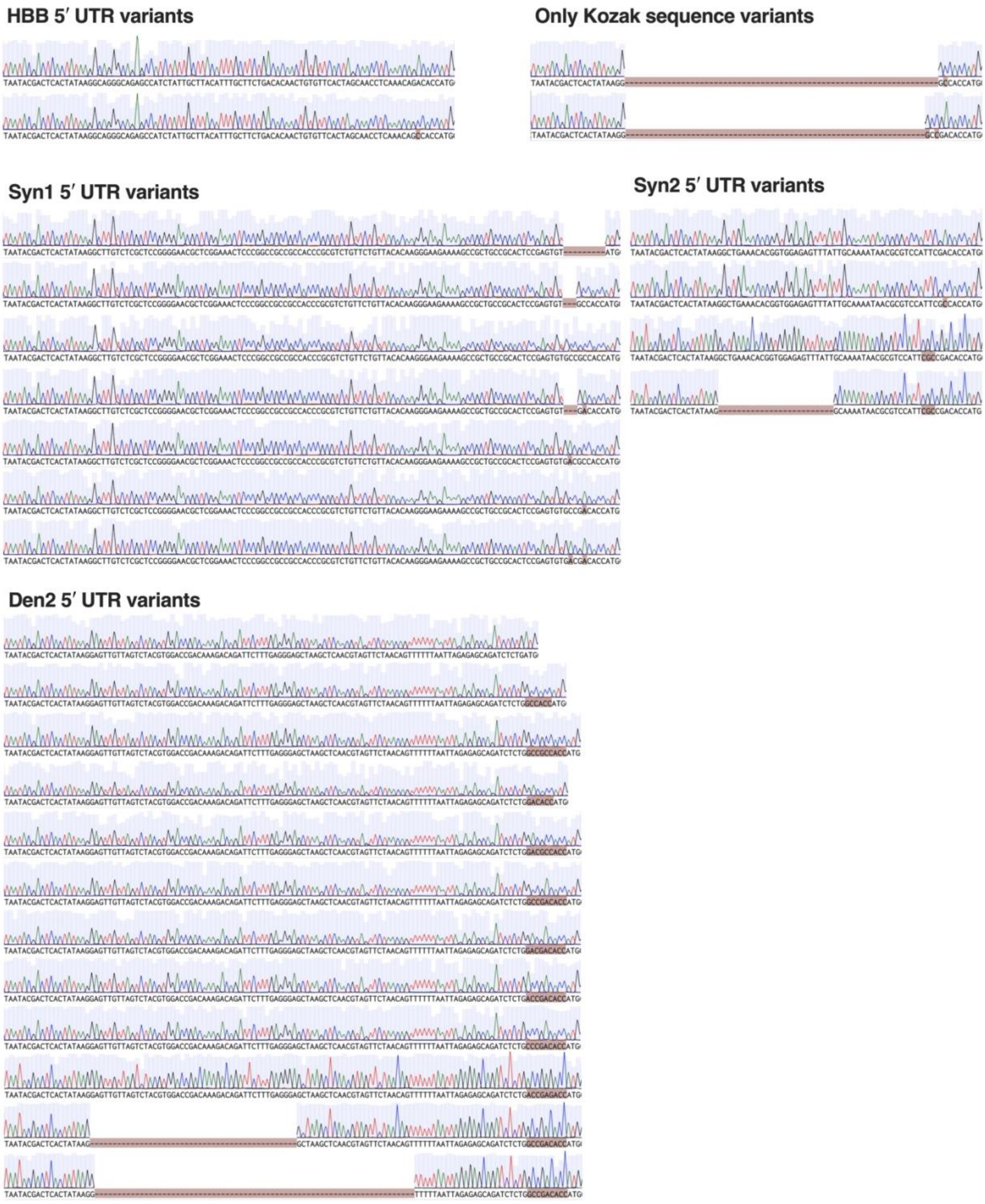


**Figure S11. Sanger sequencing reads of all 5′ UTR and Kozak variants used in this study.** BaseClear (Leiden, The Netherlands) performed Sanger sequencing on purified plasmids. Sequencing reads were aligned to the expected sequences in Benchling containing the original Kozak sequence of the respective full-length 5′ UTRs. Since Den2 originally lacks a 5′ UTR, Kozak sequences appear as gaps/mutations.

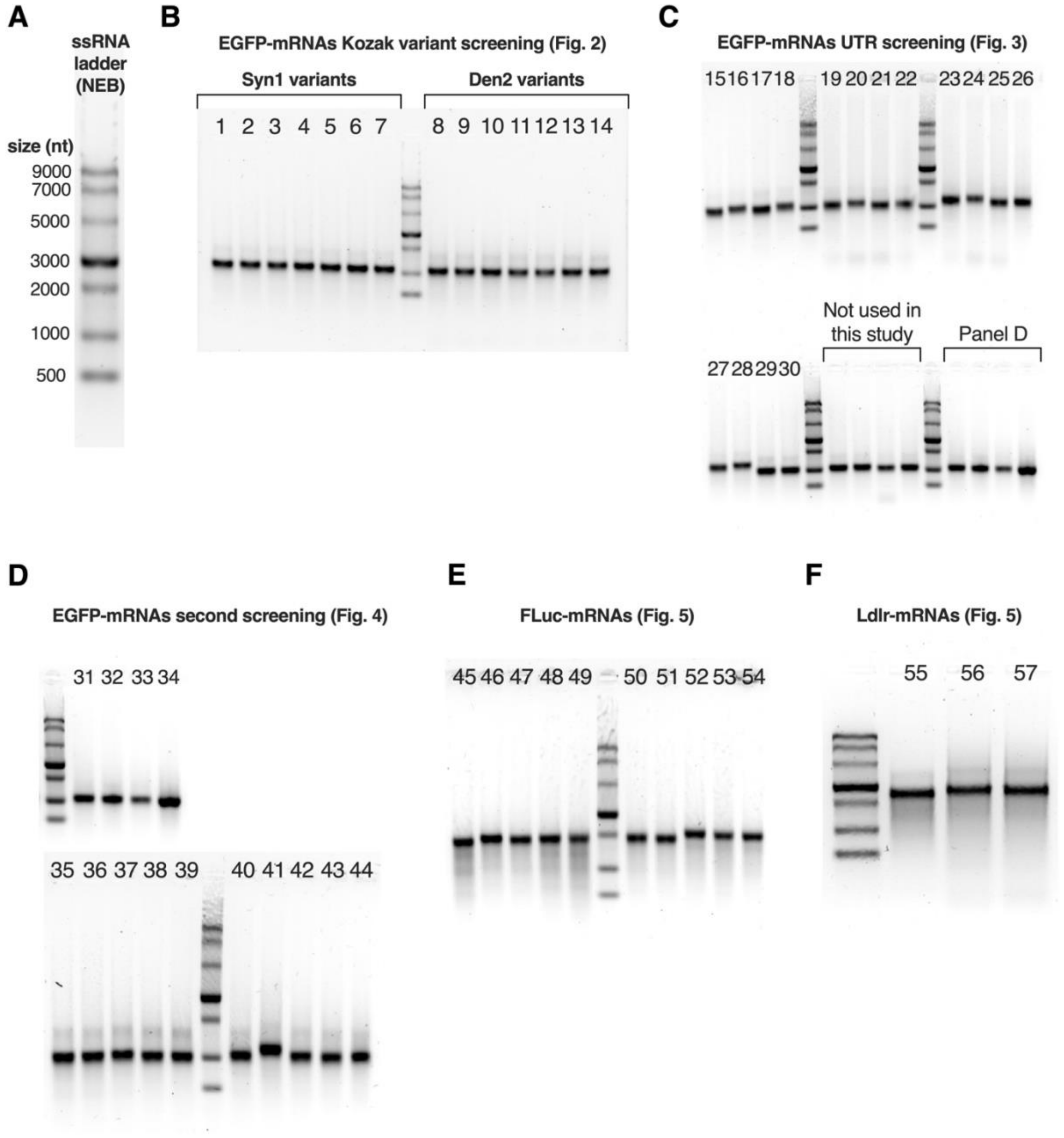


**Figure S12. Agarose gels after electrophoresis of all IVT mRNAs used in this study. A)** ssRNA ladder (NEB) used as a reference on all gels, with fragment sizes indicated from 500–9000 nucleotides. **B)** EGFP-mRNAs carrying Kozak sequence variants (Fig. 2). **C)** EGFP-mRNAs carrying UTR combinations used in combinatorial optimization (Fig. 3). **D)** All combined (truncated) UTR and Kozak variants used in the second screening (Fig. 4). **E)** FLuc-mRNAs used for expression analysis in *Ldlr* $^{-/-}$ mice. **F)** Ldlr-mRNAs used for Ldlr replacement in *Ldlr* $^{-/-}$ mice. All mRNAs are labeled with a number corresponding to the number in Table S4, which shows formulation characterization data.
